# Domain-specific decoupling of co-chaperone and ligase functions in *STUB1* underlies biochemical and clinical heterogeneity in SCA48

**DOI:** 10.1101/2025.11.26.690709

**Authors:** Selin Altinok, Ethan Paulakonis, Michael F Almeida, Isaac Hwang, Rebekah Sanchez-Hodge, Christina D’Ovidio, Georgia Roper, Grant Irons, Elena Vargas, Ivy Peng, Morcos Saeed, Aliyaa Pathan, Sarah M Ronnebaum, Change-he Shi, Mark J. Ranek, K Matthew Scaglione, Richard C. Page, Nicholas G Brown, Jonathan C Schisler

## Abstract

The carboxyl terminus of HSC70-interacting protein (CHIP, encoded by *STUB1*) integrates co-chaperone and E3 ubiquitin ligase activities to maintain proteostasis. Heterozygous *STUB1* mutations cause the dominant cerebellar ataxia SCA48 through incompletely understood mechanisms. We characterized 13 SCA48-associated variants in the TPR and U-box domains using recombinant protein and cellular assays. TPR mutations retained intrinsic ligase activity but lost HSC70 binding, substrate ubiquitination efficiency, and protein stability. Conversely, U-box mutations abolished ligase function, induced aberrant high-molecular-weight oligomerization, and frequently elevated steady-state CHIP levels while only partially impairing co-chaperone activity. Many variants exhibited temperature-sensitive defects and stress-induced nuclear mislocalization. Principal component analysis revealed robust domain-specific biochemical clustering. RNA-seq following *STUB1* knockdown demonstrated preserved HSF1-dependent transactivation but a loss of CHIP-dependent amplification of ubiquitination, chaperone, and transcriptional pathways under heat stress. Meta-analysis of 87 SCA48 patients linked TPR-like biochemical profiles to upper motor neuron involvement and U-box profiles to prominent dysarthria. Collectively, SCA48 mutations decouple CHIP’s dual functions in a domain-dependent manner, exerting dominant-negative or gain-of-toxic effects that drive the observed clinical heterogeneity. These findings establish a direct biochemical–clinical correlation in SCA48 and provide a framework for domain-targeted therapeutic strategies exploiting residual ligase or chaperone activity.

## Introduction

Spinocerebellar ataxias (SCAs) are a group of hereditary ataxias characterized by progressive gait incoordination and often linked to poor coordination of the hands, speech, and eye movements. Although the genetic causes involve mutations in various proteins, a common feature of SCAs is the inability to maintain proper protein homeostasis under stress. Therefore, strong protein quality control (PQC) responses and their machinery are vital for cellular function [1].

The gene *STUB1* (*STIP1* homology and U-box containing protein 1) encodes the PQC protein known as CHIP (Carboxyl terminus of the Hsc70-interacting protein). Importantly, mutations in *STUB1* can cause a wide range of clinical symptoms, including classic ataxia, developmental delay, cognitive decline, accelerated aging, and hypogonadism. The first disease-related mutation in *STUB1* was found in patients with a new recessive form of spinocerebellar ataxia, SCAR16 [2,3]. In 2018, a heterozygous *STUB1* mutation was linked to a new autosomal dominant spinocerebellar ataxia, SCA48 [4]. Additionally, *STUB1* mutations have been shown to modify other ataxias, such as SCA8 and SCA17 [5,6]. These findings collectively highlight a unique role for *STUB1* as both a driver and modifier of SCAs, emphasizing its importance in the development of these disorders.

CHIP plays a dual role in the PQC pathway as a co-chaperone with HSPs and as a ubiquitin ligase, helping proteins fold and be targeted for degradation, respectively [7–10]. CHIP’s multiple enzymatic functions highlight its importance in the cellular stress response, especially in neurodegenerative diseases [11–15]. Additionally, understanding how these mutations disable CHIP’s various functions is crucial, as it points to CHIP’s potential as a therapeutic target because of its ability to ubiquitinate and target disease-related proteins for proteasomal degradation [16].

CHIP uses TPR (tetratricopeptide) and U-box domains to perform its enzymatic functions. In brief, the N-terminal TPR domain binds to chaperones, such as HSP(C)70, for protein refolding and recruits substrates for ubiquitination. CHIP’s inherent ubiquitin (Ub) ligase activity is enabled by the U-box domain, which binds and activates E2 Ub-conjugating enzymes, like UBE2D2. The co-recruitment of substrates to its TPR and to the E2∼Ub (“∼” indicates a transient thioester intermediate) to its U-box facilitates polyubiquitination. Synthetic mutations in each domain, such as TPR (K30A) and U-box (H260Q), have been used previously to investigate the functions of each domain. Besides its enzymatic roles, CHIP naturally forms stable dimers and higher-order oligomers of repeating dimers, like tetramers and hexamers, for the heat shock response [17–19]. Prior studies have shown that SCAR16 disease-related mutations destabilize CHIP, resulting in a loss of co-chaperone or ubiquitin ligase activity [20]. Additionally, a well-characterized SCA48 mutation in the TPR domain (A52G) was defective in recruiting Hsp70 [21]. However, other SCA48-linked mutations occur in both the TPR and U-box domains of CHIP [22,23], which may lead to a loss of CHIP’s ubiquitin ligase activity and a gain of toxic function, potentially related to disease symptoms.

Despite the critical role of CHIP in maintaining protein homeostasis and its involvement in various neurodegenerative diseases, there are currently no effective therapies targeting *STUB1* mutations. Therefore, understanding the molecular mechanisms behind these mutations is essential for developing targeted treatments. Ultimately, our findings reveal that mutations cause a complex range of changes in protein stability, oligomerization, and enzymatic activity, depending on their location. Furthermore, these biochemical changes are associated with clinical manifestations in SCA48 patients.

## Results

### SCA48 mutations have different effects on CHIP protein expression levels

SCA48 mutations encompass both single amino acid substitutions and protein domain deletions. Therefore, we chose a subset of 13 SCA48-associated mutations within the TPR and U-box domains for detailed analysis, including seven missense mutations, five frameshift mutations, and one nonsense mutation [22,23]. We also examined a *STUB1* mutation in the start codon, c.3G>A, which results in a truncated TPR domain due to an in-frame alternative translation start site within the second exon (denoted as ΔStartCodon) [24] (**Figure 1A**).

**Figure 1:**
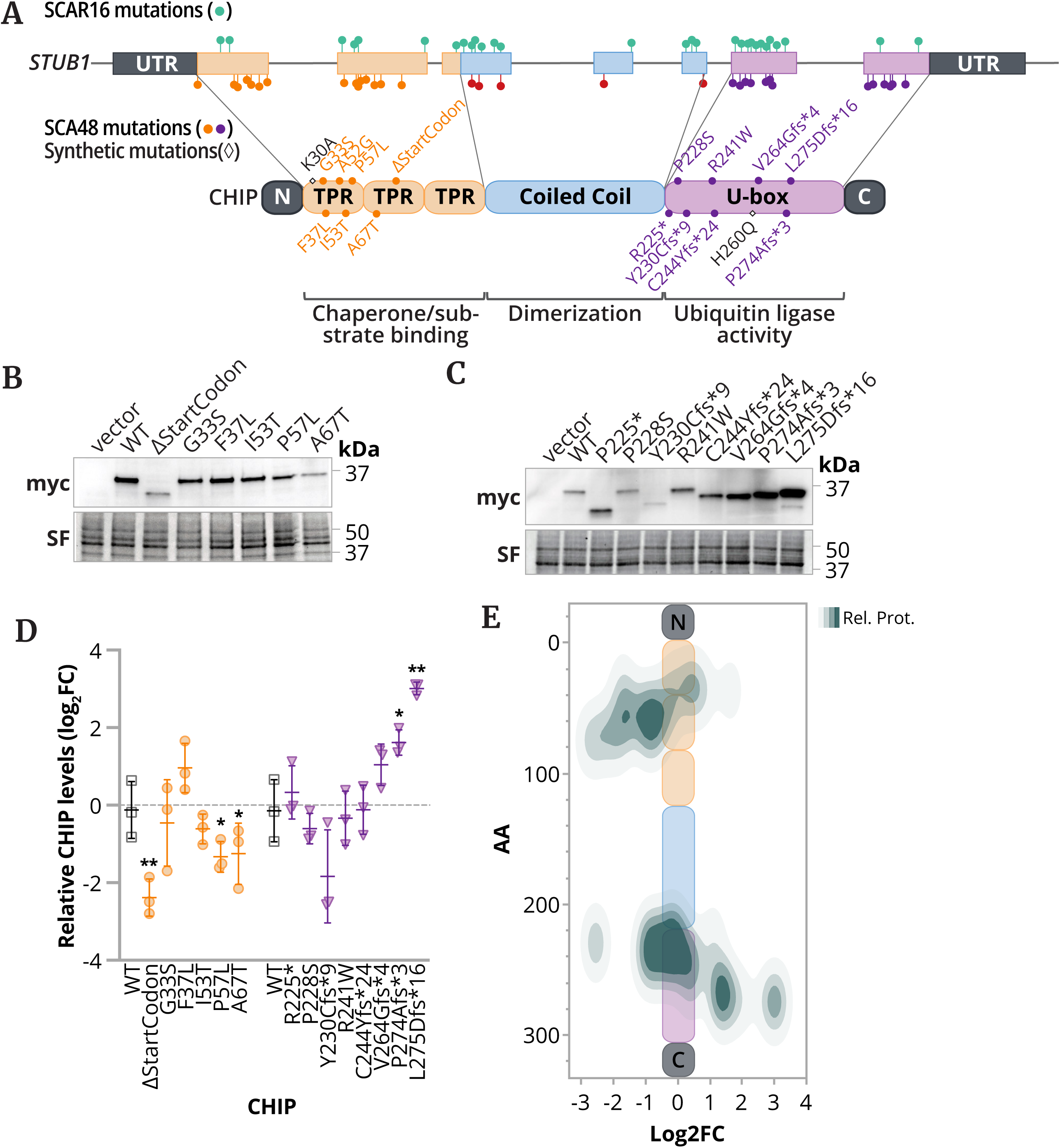
Distinct CHIP protein expression patterns of SCA48-linked mutations. (A) Gene diagram showing *STUB1* mutations associated with recessive SCAR16 (green, upper) and dominant SCA48 (orange, red, and purple, lower), and the resulting location in the encoded CHIP protein, highlighting selected SCA48-linked mutations in the TPR and U-box domains, along with synthetic mutations (K30A and H260Q). (**B & C**) Representative SDS-PAGE immunoblots of steady-state protein levels of the indicated myc-tagged CHIP protein transiently expressed in COS-7 with stain-free (SF) levels of total protein. (**D**) Densitometry analysis of normalized CHIP protein levels represented by the log2 fold change, summarized by a dot plot, and represented by the mean ± SD (n = 3). Changes to steady-state levels were determined via one-sample t-test, * and ** indicating P < 0.05 and < 0.01, respectively. (**E**) Density heatmap plotting protein levels (Rel. Prot.) relative to WT CHIP and the amino acid (AA) position of the mutation.

First, we evaluated CHIP’s steady-state levels in a cell culture system by transiently transfecting COS-7 cells with constructs containing different CHIP mutations. By assessing protein levels through immunoblotting, we observed that TPR domain mutants either showed no change or had reduced protein expression. For example, the ΔStartCodon, which truncates the TPR domain, was markedly decreased, likely due to protein misfolding. Conversely, U-box mutants displayed varying protein expression levels, with some mutations leading to decreased levels and others to increased levels (**Figure 1B-D**). This wide range of protein expression suggests that distinguishing between changes in protein function and changes in protein levels is difficult solely in cell-based systems. However, plotting the relative change in expression along the primary structure of CHIP reinforced this trend: lower expression in TPR mutants and higher expression in U-box mutants, suggesting that the stability contributed by these domains may differ in cells (**Figure 1E**).

### Recombinant systems exhibit notable differences in protein stability and oligomerization

The varied protein levels across the SCA48-harboring mutations could result from changes in CHIP’s intrinsic properties, such as stability, oligomerization, enzymatic activity, or a combination of these factors. For instance, defects in CHIP’s ubiquitination activity might stabilize the steady-state protein levels by decreasing autoubiquitination and degradation. To investigate these fundamental characteristics in an isolated, cell-free system, we purified recombinant forms of both wild-type and CHIP disease variants to perform a series of *in vitro* assays.

Given the dominant inheritance pattern of SCA48, we hypothesized that SCA48 mutations would be as stable or more stable than wild-type CHIP. Therefore, to initially assess protein stability, we conducted differential scanning fluorimetry on recombinant CHIP proteins. These results did not show a consistent trend in thermal stability among SCA48 proteins. Instead, we observed varied effects across different variants (**Table 1**). For example, while P57L and Y230Cfs*9 showed increased stability, the A67T and R241W mutations significantly lowered the melting temperature of CHIP, indicating stabilization and destabilization, respectively. Several variants with lower melting temperatures, such as G33S, II53T, A67T, and R241W, also exhibited predicted steric clashes when modeled on the CHIP dimeric structure (**Supplementary Figure 1**). These changes in protein stability may help explain the lower steady-state levels observed for some CHIP variants.

**Table 1.**
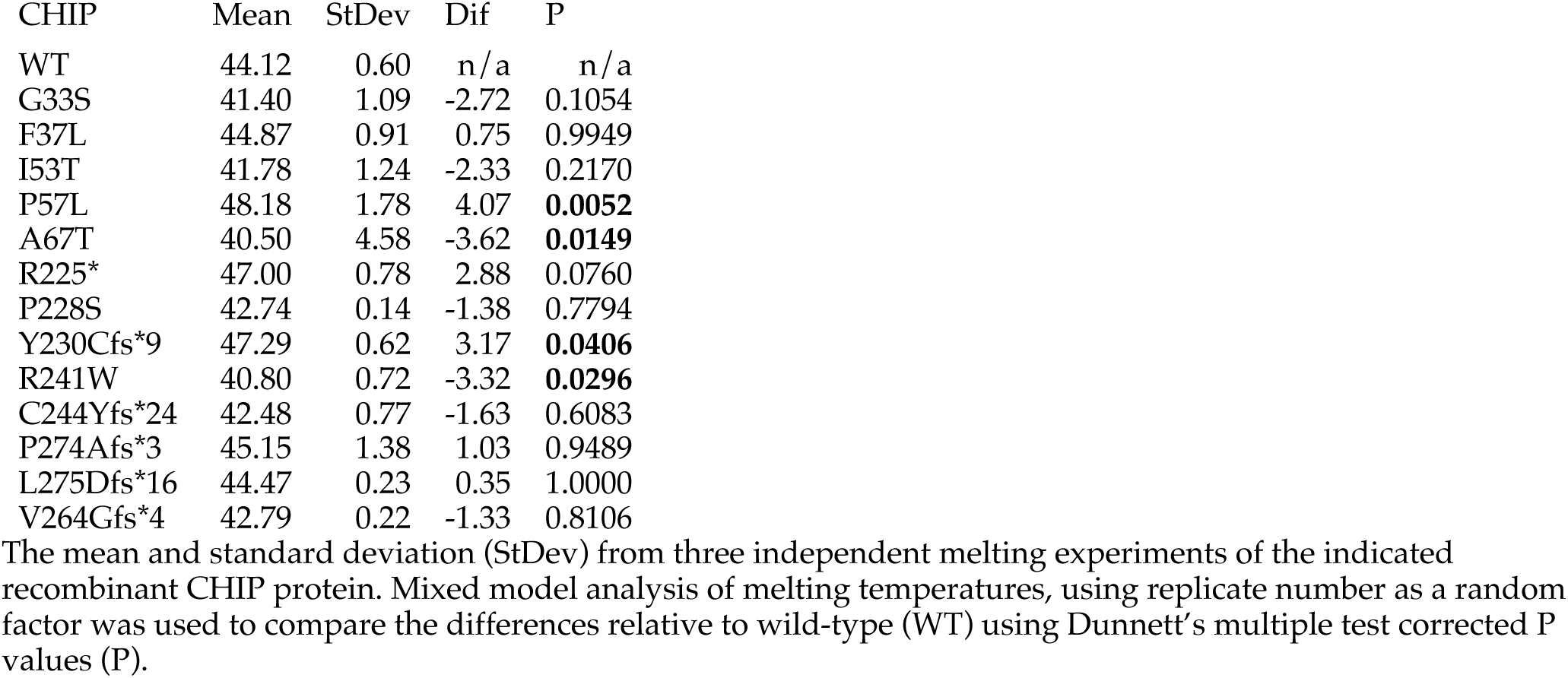
Melting temperatures of SCA48-linked CHIP mutations.

Next, we used mass photometry (MP) to monitor the oligomerization of CHIP wild type and variants, as a possible gain of toxic function might be due to an increased propensity for protein aggregation. We observed distinct peaks at molecular weights corresponding to dimer, tetramer, hexamer, and octamer with CHIP WT (**Figure 2A-B**). All TPR domain mutants showed clear peaks matching all four oligomeric states (**Figure 2A-B & Supplementary Figure 2A-B**). The R241W variant was the only U-box domain mutant that oligomerized similarly to CHIP WT (**Supplementary Figure 2A-B**). Interestingly, about 80% of R225* and Y230Cfs*9 proteins existed as dimers, lacking clear higher-order states such as hexamer and octamer (**Supplementary Figure 1C-D**). Conversely, the other U-box mutants mainly formed large, non-distinct high-order oligomers over 200 kDa, with minimal dimer presence (**Supplementary Figure 2E-F**). Based on their MP profiles, we used principal component analysis to categorized SCA48-associated CHIP mutations into three groups (**Figure 2C**) : (1) dimer repeating units (DR), which show distinct oligomeric states at dimer, tetramer, hexamer, and octamer; (2) dimer-only (D), for mutations with approximately 80% dimer formation and no clear high oligomeric state; and (3) non-discrete high molecular weight (HMW), representing mutants that produce very low levels of distinct oligomers and are mostly found as non-distinct high-order oligomers (**Table 2**).

**Figure 2.**
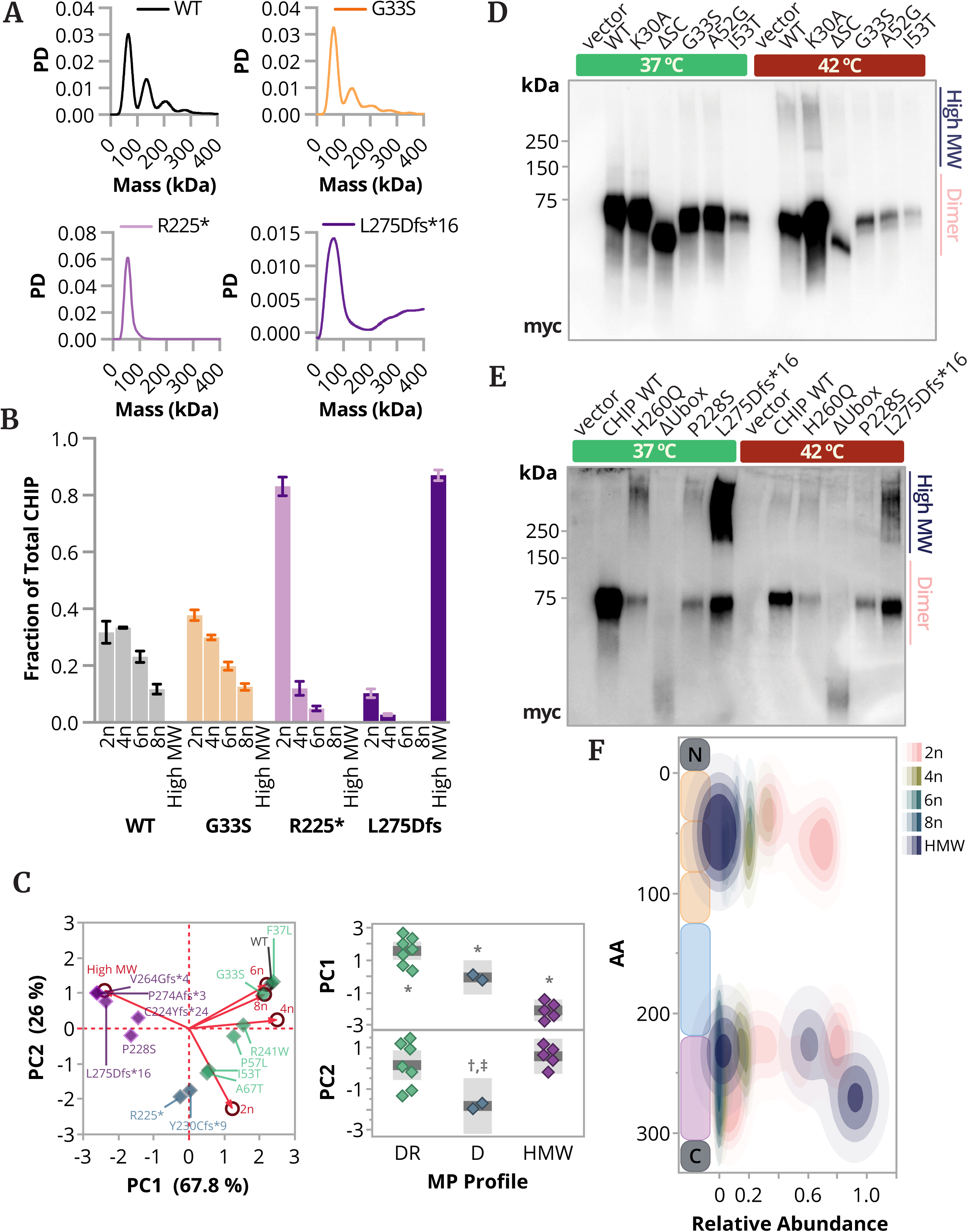
Effect of SCA48-associated mutations on CHIP oligomeric state. (**A**) Probability distribution (PD) versus molecular mass (kDa), determined by mass photometry, of WT CHIP and three SCA48 mutant proteins that represent the main patterns of CHIP oligomerization. Full plots and additional data are present in Supplementary Figure 2. (**B**) The proportions of CHIP oligomeric states, dimer (2n), tetramer (4n), hexamer (6n), octamer (8n), and high molecular weight (High MW), are represented by a bar graph and summarized by the mean ± SD (n = 3). (C) Oligomeric variation of CHIP proteins represented by (left) scatterplot of the first two principal components (PC) with the loadings (◯) represented by red arrows and proteins (◇) colored by mass photometry (MP) profile, and (right) dot plot of MP profiles across PC1 (top) and PC2 (bottom). Oneway ANOVA of PC1 and PC2, F(2,11) = 43, P < 0.0001, and F(2,11) = 5.7, P = 0.0201. Tukey posttest: * P < 0.05 of all pairwise comparisons, † P < 0.05 D vs. DR, and ‡ P < 0.05 D vs. HMW. The dark line represents the mean, and the shaded region represents the 95% confidence interval (**D & E**). Immunoblot analysis of CHIP oligomerization in COS-7 cells transfected with the indicated plasmids using native blue PAGE, under either 37 °C or 42 °C conditions, with dimer and high MW species indicated. ΔStartCodon is abbreviated as ΔSC. (**F**) Density heatmap showing amino acid (AA) positions and the resulting abundance of CHIP oligomeric species.

**Table 2.**
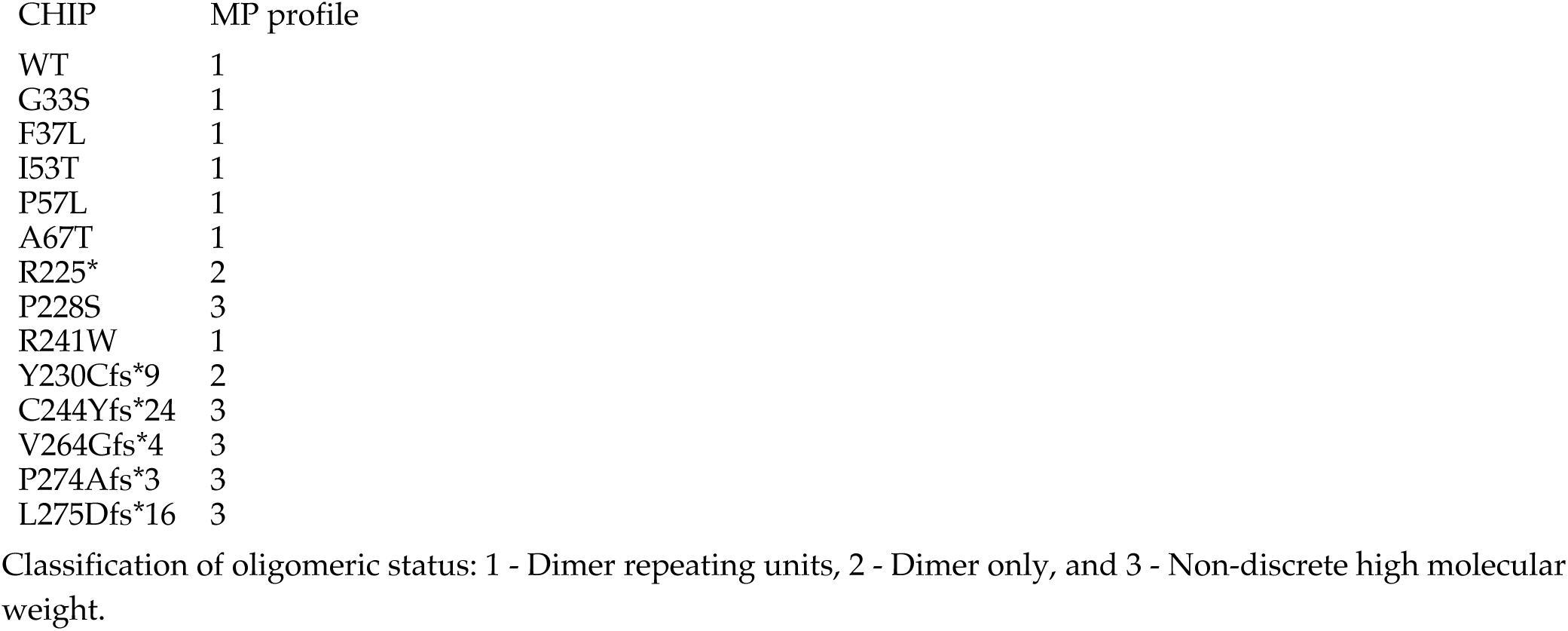
MP profile of CHIP mutations.

| CHIP | MP profile |
| --- | --- |
| WT | 1 |
| G33S | 1 |
| F37L | 1 |
| I53T | 1 |
| P57L | 1 |
| A67T | 1 |
| R225* | 2 |
| P228S | 3 |
| R241W | 1 |
| Y230Cfs*9 | 2 |
| C244Yfs*24 | 3 |
| V264Gfs*4 | 3 |
| P274Afs*3 | 3 |
| L275Dfs*16 | 3 |

Classification of oligomeric status: 1 - Dimer repeating units, 2 - Dimer only, and 3 - Non-discrete high molecular weight.

The oligomeric state of proteins often influences their function, especially during cellular stress. For example, CHIP forms high-order oligomers after heat shock, and its chaperone activity increases with heat, indicating that mutations affecting oligomerization can impact CHIP’s function [17]. Therefore, we aimed to identify changes in the oligomeric state of CHIP wild type and those with SCA48 mutations in a cellular system after heat shock. COS-7 cells were transfected with CHIP wild-type and mutant constructs and lysed under non-denaturing conditions to preserve native structure and interactions. CHIP oligomerization was analyzed using Blue Native-PAGE followed by immunoblotting. At physiological temperature, CHIP WT and all TPR mutants mainly appeared as dimers (**Figure 2D**), as observed in MP (**Figure 2A-B**). When subjected to heat shock at 42 °C, high-molecular-weight species of CHIP WT and K30A formed, while disease-related TPR mutants failed to form high-order oligomers (**Figure 2D**). Consistent with earlier MP data, U-box mutants appeared as high-order oligomers larger than 200 kDa in COS-7 cells at physiological temperature (**Figure 2E**). However, after heat shock, there was a reduction in high-order oligomers of U-box mutated proteins (**Figure 2E**). Mapping MP-derived oligomeric status of the mutations across the primary structure of CHIP again reinforces domain-specific trends in oligomerization (**Figure 2F**).

Along with MP data, these results show consistent changes in the oligomeric state of CHIP, which may alter its enzymatic and co-chaperone activities. In its monomer form, CHIP is considered inactive [25]. Asymmetric dimerization of two protomers exposes the E2 binding region of the U-box domain, making it accessible for substrate ubiquitination [25,26]. Since these data indicate differences in CHIP’s biophysical properties, any changes in oligomerization and protein-protein interactions could also affect its enzymatic ubiquitin ligase and co-chaperone functions.

### SCA48 mutations in the TPR and U-box differentially impact CHIP-dependent ubiquitination activity

Since its discovery, the ubiquitin (Ub) ligase activity of CHIP has been widely studied [8,9,27–29]. CHIP facilitates the ubiquitination of molecular chaperones such as HSC70 and their bound substrates. Specifically, CHIP recruits co-chaperones and conjugated E2∼Ub through its TPR and U-box domains, respectively [9,10]. To assess the substrate ubiquitination activity of the SCA48 variants, we conducted CHIP-dependent ubiquitination assays using fluorescently labeled HSC70 (**Figure 3A-B**). Overall, we observed reduced HSC70 ubiquitination across all SCA48 variants, except A67T and R241W.

**Figure 3.**
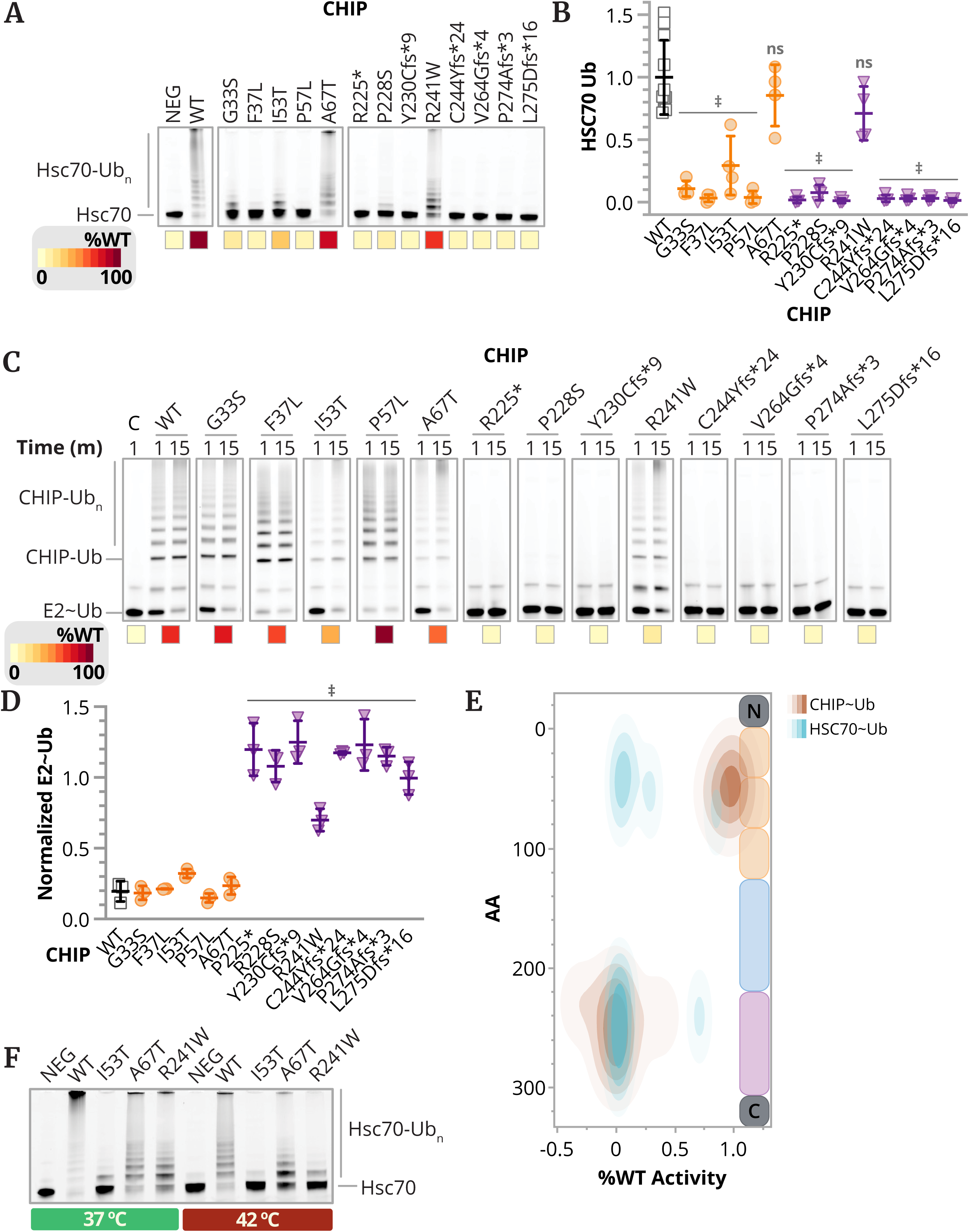
Distinct effects of SCA48 mutations on CHIP-dependent ubiquitination activity. (**A**) Representative gel images of CHIP-dependent ubiquitination of fluorescently labeled HSC70, with increasing ubiquitination denoted by HSC70-Ub^n^ and summarized by a cell plot of total HSC70 ubiquitination relative to WT CHIP. (**B**) Densitometry analysis of HSC70 ubiquitination (HSC70 Ub) by each CHIP variant, represented by a dot plot and summarized by the mean ± SD (n = 4 - 11), was analyzed via one-way ANOVA (F(13, 49) = 26.7, P = 0.0303) and Dunnett’s multiple comparison test, ‡ and ns, indicate P < 0.0001 and > 0.05, respectively. (**C**) Representative gel images of CHIP autoubiquitination assay using fluorescently labelled ubiquitin (Ub) and summarized by a cell plot of autoubiquitination levels of each CHIP protein relative to WT CHIP. (**D**) Densitometry analysis of E2∼Ub levels from each condition (lower levels equate to higher autoubiquitination), represented by a dot plot and summarized by the mean ± SD (n = 3), was analyzed via one-way ANOVA (F(13, 28) = 67, P < 0.0001) and Dunnett’s multiple comparison test, ‡ P < 0.0001. (**F**) Representative gel images of CHIP-dependent ubiquitination of fluorescently labeled HSC70, with increasing ubiquitination denoted by HSC70-Ub^n^ at 37 °C and 42 °C compared to **C**. (**E**) Density heatmap of CHIP autoubiquitination (CHIP-Ub) and HSC70 ubiquitination relative to WT CHIP conditions and the amino acid position of the mutation.

Since CHIP’s ubiquitination activity toward HSC70 relies on a functional U-box domain and binding to the TPR domain, we conducted *in vitro* autoubiquitination assays using a fluorescently labeled Ub to directly assess the intrinsic Ub ligase functions of the CHIP variants. As expected, mutations in the TPR domain largely did not impair its intrinsic ligase activity, as shown by the increase in CHIP-Ub^n^ bands and the decrease in ubiquitin-conjugated E2 (E2∼Ub) bands (**Figure 3C-D**). In contrast, U-box domain mutations failed to exhibit autoubiquitination activity, except for some activity observed with R241W, suggesting an inactive U-box domain. These findings demonstrate that mutations in the TPR and U-box domains primarily reduce substrate ubiquitination activity through two mechanisms: failure to recruit substrates or inactivation of the U-box domain (**Figure 3E**).

Three exceptions to this concept were I53T, A67T, and R241W. These enzymes showed a higher level of ubiquitination than the other mutants tested at 25 °C. However, CHIP activity is crucial during heat stress, and these mutations had among the lowest melting temperatures, suggesting that their ubiquitination activity might decline at higher temperatures. When their ubiquitination activity was tested at physiological temperature (37 °C) or heat-stress temperature (42 °C), CHIP-dependent HSC70 ubiquitination was notably reduced at the higher temperatures compared to wild-type CHIP, indicating that these variants would likely be defective in ubiquitination within cells (**Figure 3F**).

### SCA48 TPR mutations result in defective co-chaperone binding and activity

The observation that TPR substitutions lead to a loss of HSC70 ubiquitination but maintain its intrinsic ligase function suggests that these variants might also be impaired in their HSP70-dependent co-chaperone activity. Conversely, we hypothesized that U-box mutations would preserve their co-chaperone function, similar to what has been observed with SCAR16 U-box mutations [19,20]. To begin assessing CHIP-HSP70 binding, we performed co-immunoprecipitation on lysates from COS-7 cells transfected with HSP70 and a select set of CHIP variants. Overall, less HSP70 co-precipitated with CHIP containing TPR mutations, including the previously identified SCA48 mutant A52G [21]. In contrast, U-box mutants efficiently co-precipitated HSP70 (**Figure 4A**). These results further indicate that the reduced ubiquitination of HSC70 in CHIP variants with TPR substitutions results from a deficiency in substrate recruitment.

**Figure 4.**
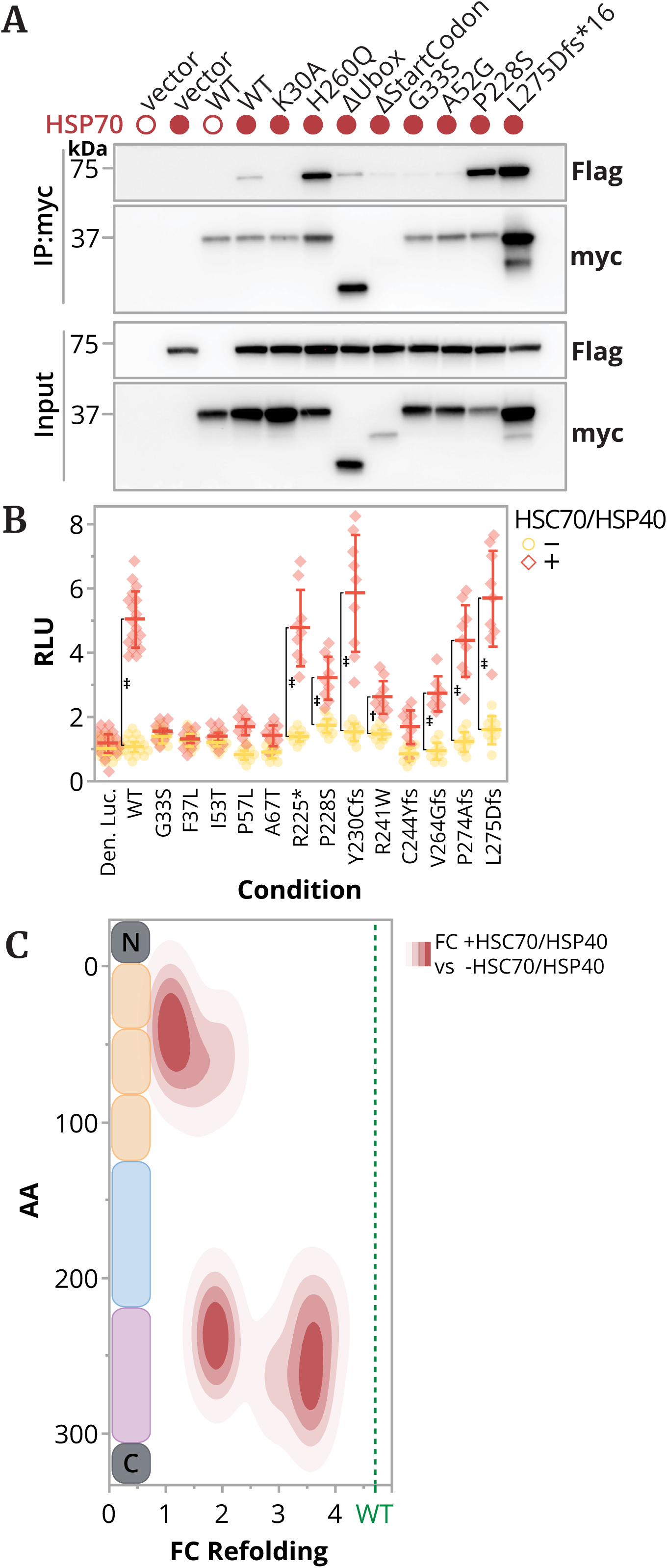
SCA48 mutations differentially affect co-chaperone activity. (**A**) Co-immunoprecipitation (IP) of CHIP and HSP70, along with input levels of both transgenes visualized by immunoblot, from COS-7 cells co-expressing the specified HSP70 and CHIP variants. (**B**) Co-chaperone refolding activity was assessed by measuring relative luciferase units (RLU) in heat-denatured luciferase refolding assays, with recombinant CHIP protein present (+) or absent (−) alongside HSC70 and HSP40. A two-way mixed model ANOVA of RLU (considering HSC70/HSP40 and CHIP), with replicate (n = 11) and day (n = 3) as random factors, resulted in F(14, 14) = 50, P < 0.0001 for the interaction term. Tukey post-tests indicated † and ‡, P < 0.01 and P < 0.0001, respectively, when comparing conditions with and without HSC70/HSP40. (**C**) Density heatmap of co-chaperone activity, summarized by fold change in luciferase refolding (FC refolding) in the presence of HSC70/HSP40 and the mutation’s amino acid position. Wild-type (WT) CHIP activity is indicated by the dashed line.

The interaction between HSC(P)70 and the CHIP TPR domain is also essential for protein refolding and can be directly tested by measuring the refolding of heat-denatured luciferase in the presence of HSC70 and HSP40. While luciferase refolding required both CHIP, HSC70, and HSP40, SCA48-associated substitutions in the TPR domain led to decreased luciferase refolding (**Figure 4B-C**) [17,19]. In contrast, the U-box variants showed different levels of luciferase refolding activity. Some frameshift variants and R225* maintained activity similar to CHIP WT, while P228S, R241W, and C244Yfs*24 exhibited reduced co-chaperone activity (**Figure 4B-C**). Other than R241W, which is temperature sensitive, P228S and C244Yfs*24s were classified as forming non-discrete high molecular weight variants based on MP (**Supplementary Figure 2E**), suggesting that U-box mutations can impact more than CHIP’s established ubiquitin ligase activity.

### Domain-specific biochemical and structural divergence in SCA48 mutations

To characterize the biochemical and structural diversity of SCA48 mutations, we examined 13 mutations in the TPR and U-box domains of CHIP, combining multivariate, molecular, and domain-specific summaries (**Figure 5**).

**Figure 5.**
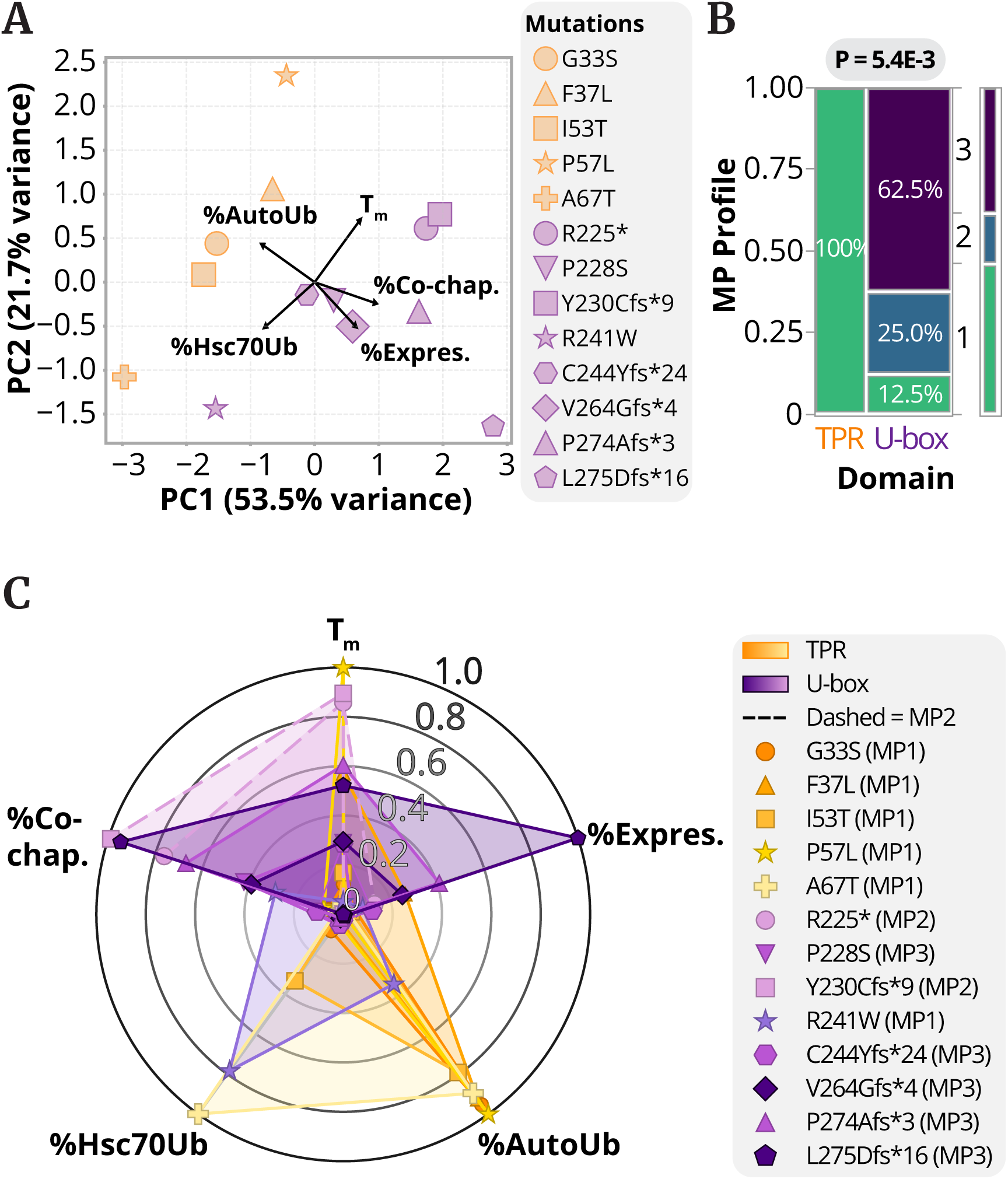
Integrated biochemical and structural profiling reveals domain-dependent mechanisms of SCA48-associated CHIP dysfunction. (**A**) Principal component analysis (PCA) of normalized biochemical metrics: thermal stability (T^m^); steady-state expression (% Expres.); auto ubiquitination (%AutoUb); HSC70-dependent ubiquitination (%HSC70Ub); and co-chaperone activity (%Co-Chap), represented by a scatterplot. The first two principal components, PC1 (53.5%) and PC2 (21.7%), together explain 75% of the total variance, and domain-dependent separation on PC1 is confirmed via t-test (t(11) = 3.3, P = 0.0069). (**B**) Contingency analysis of mass photometry (MP) profile distribution by domain reveals unequal distributions with P = .0054 via Fisher’s Exact Test. (**C**) Radar plot of the indicated biochemical quantitative metrics (normalized to WT CHIP = 1.0) for 13 *STUB1* mutations.

#### Principal component analysis uncovers domain-specific biochemical clustering

We performed a principal component analysis (PCA) on our normalized biochemical metrics, including thermal stability, expression levels, auto-ubiquitination, HSC70 ubiquitination, and co-chaperoning, to explore multidimensional relationships (**Figure 5A & Supplementary Table 1**). The first two principal components (PC1 and PC2) explained 75% of the total variance, with PC1 accounting for 53% and PC2 for 22%. The PCA revealed distinct clusters: TPR mutations (G33S, F37L, I53T, P57L) mostly grouped in the quadrant characterized by autoubiquitination loading, suggesting that autoubiquitination is a key feature of TPR mutations. A67T (TPR) and R241W (U-box) diverged, with %HSC70 ubiquitination as the main driver, indicating a unique substrate ubiquitination profile. U-box mutations R225* and Y230Cfs\**9* are mainly affected by Tm, while the other U-box mutations (P228S, C244Yfs*24, V264Gfs*4, P274Afs*3, and L275Dfs*16) clustered with co-chaperone activity and expression loadings. A t-test on PC1 scores confirmed significant separation between domains (t(11) = 3.3, p = 0.0069), supporting the biochemical distinction between TPR and U-box mutations.

#### Mass photometry identifies domain-specific molecular assemblies

Structural insights from mass photometry (MP) profiling highlighted significant domain-dependent differences in molecular assembly (**Figure 5B**). All TPR mutations (G33S, F37L, I53T, P57L, A67T) exhibited an MP profile 1, characterized by dimer repeat units, whereas U-box mutations were predominantly MP profile 3, indicative of high molecular weight aggregates (R225*, P228S, Y230Cfs*9, C244Yfs*24, V264Gfs*4, P274Afs*3, L275Dfs*16), with R241W and P228S as MP profile 2 (dimer only). A contingency analysis confirmed this distribution (p = 5.4 × 10⁻³), suggesting that aggregation propensity may differentiate TPR and U-box pathobiology, potentially influencing cellular localization and function.

#### Integrating domain-specific biochemical signatures

A radar plot visualized the normalized biochemical metrics across all 13 mutations, highlighting domain-specific patterns (**Figure 5C & Supplementary Table 1**). TPR mutations maintained intact U-box function, consistent with their PCA clustering and MP Profile 1 dimerization. Conversely, U-box mutations showed increased steady-state expression levels and co-chaperone activity, aligning with their aggregate-prone MP Profile 3 and PCA distribution. Overall, thermal stability and steady-state expression levels exhibited the most mutation-specific variability within each domain, providing a comprehensive overview that connects structural and functional data to downstream cellular and clinical outcomes.

### Mechanistic insights into CHIP dysfunction in SCA48

The radar plot (**Figure 5C & Supplementary Table 1**) summarizes our biochemical profiling and reveals distinct domain-specific signatures among SCA48 mutations in the *STUB1*/CHIP protein: TPR variants, such as G33S, maintain ubiquitination activity but have reduced chaperone function and expression, while U-box variants, including P228S and L275Dfs16, show preserved chaperone interactions but nearly no ligase activity. These contrasting profiles suggest a decoupling of substrate recognition and degradation, which may contribute to proteostatic imbalance in affected cells. To examine these domain-specific effects, we performed cell-based assays on these selected mutants: G33S (a TPR missense, MP1), P228S (a U-box missense, MP3), and L275Dfs16 (a U-box frameshift, MP3), focusing on protein stability, post-translational modifications, stress responses, subcellular localization, and transcriptional regulation (**Figures 6 & 7**).

**Figure 6.**
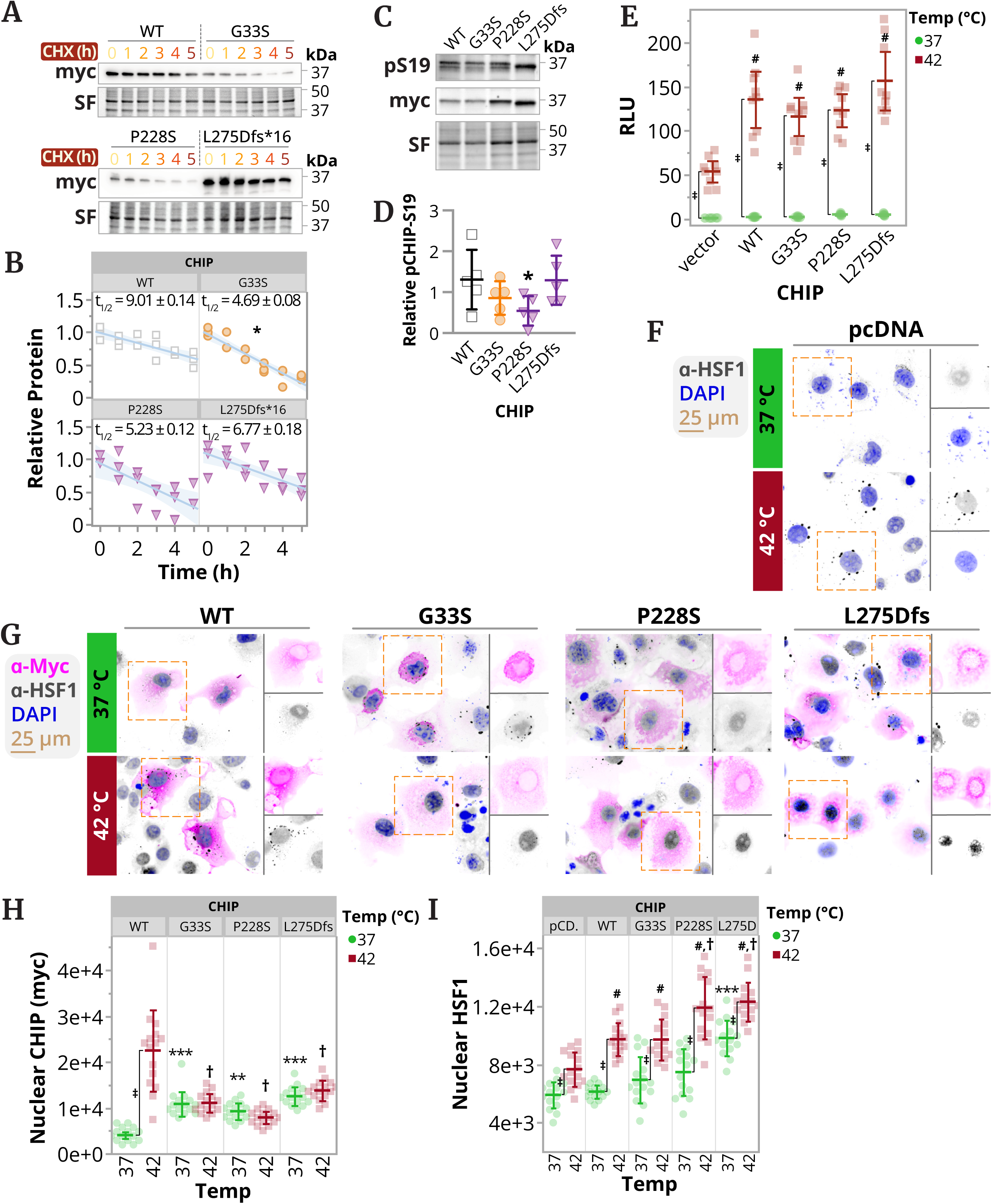
Impact of SCA48 mutations on CHIP stability, phosphorylation, and heat shock response regulation. (**A**) Representative immunoblot of cycloheximide (CHX) chase assays detecting the myc-tag of exogenous CHIP vectors with stain-free (SF) loading confirmation, and (**B**) densitometry analysis represented by scatterplot. Half-lives were calculated from robust linear regression as t^1/2^ = ln 2 * (∣slope∣^-1^); * P = 0.00015 with Dunnett’s adjusted post-test. (**C**) Representative immunoblot of indicated CHIP proteins phosphorylated at S19, total exogenous CHIP (myc), and stain-free loading confirmation. (**D**) Densitometry analysis of relative pS19 represented by a dot plot, summarized by the mean ± SD (n = 5), analyzed via one-sample t-test, * P < 0.05. (**E**) HSF1 activity measured by normalized relative luciferase units (RLU) using a dual luciferase reporter assay represented by a dot plot, summarized by the mean ± SD (n = 9), and analyzed using a two-way mixed model ANOVA of RLU (main factors = temperature and CHIP), with replicate (n = 3) and day (n = 3) as random factors, resulted in F(4, 4) = 11.7, P < 0.0001 for the temperature*CHIP interaction term. Tukey post-tests indicated # and ‡ at P < 0.0001, when comparing each mutant protein to WT at 42 °C and 42 °C vs. 37 °C within each CHIP protein condition, respectively. (**F**) Representative false colored immunofluorescence composite micrographs from COS-7 cells transfected with the control vector (pcDNA) at 37 °C and 42 °C stained for HSF1 (gray) with DAPI nuclear counter stain (blue) in COS-7 cells or (**G**) additional staining for WT or the indicated mutant exogenous CHIP (myc, magenta). Dashed boxes indicate the selected cell included in the single-channel insets (**H & I**). Quantification of nuclear abundance of either CHIP (myc) or HSF1 at 37 °C and 42 °C is represented by dot plots and summarized by the mean ± SD (n = 30 cells per condition), analyzed with two-way ANOVA with temperature and CHIP variant as main factors. For nuclear CHIP, temperature, CHIP variant, and the interaction term, F(1,1) = 50, P < 0.0001; F(4,4) = 86, P < 0.0001; and F(4,4) = 49, P < 0.0001, respectively. Tukey post-test: ‡ P < 0.0001 (37 °C vs 42 °C); ** P < 0.001 and *** P < 0.0001 (vs WT at 37 °C); and † P < 0.0001 (vs WT at 42 °C). For nuclear HSF1, temperature, CHIP variant, and the interaction term, F(1,1) = 183, P < 0.0001; F(4,4) = 44, P < 0.0001; and F(4,4) = 4.2, P = 0.0028, respectively. Tukey post-test: ‡ P < 0.0001 (37 °C vs 42 °C); *** P < 0.0001 (vs WT at 37 °C); and # P < 0.001 (vs pcD. at 42 °C); and † P < 0.001 (vs WT at 42 °C).

To assess the impact on protein turnover, we conducted cycloheximide (CHX) chase assays to determine the half-lives of WT CHIP and the mutants (**Figure 6A-B**). WT CHIP demonstrated a half-life of 9.01 ± 0.14 hours, indicating high stability. In contrast, the TPR mutant G33S had a significantly shorter half-life of 4.69 ± 0.08 hours (P < 0.05, Dunnett’s test vs. WT), which corresponds with the increased turnover observed with the related TPR mutant A52G [21]. U-box mutants P228S (5.23 ± 0.12 hours) and L275Dfs*16 (6.77 ± 0.18 hours) exhibited intermediate half-lives, with no significant difference from WT (P > 0.05), suggesting a milder effect on stability.

#### Stabilizing phosphorylation of CHIP does not explain differences in half-life

Given the role of post-translational modifications in regulating CHIP stability, we examined phosphorylation at serine 19 (pS19), a modification previously linked to increased stability [30]. Western blot analysis showed P228S had lower pS19 levels compared to WT, while G33S and L275Dfs*16 maintained WT-like levels (**Figure 6C-D**). This decrease in P228S suggests a possible reduction in pS19-mediated stability; however, its half-life was similar to WT, indicating that pS19 levels alone do not determine steady-state abundance. Conversely, the significant decrease in half-life observed in G33S (**Figure 6A-B**), despite unchanged pS19, suggests other regulatory mechanisms, possibly related to its TPR domain impairment, which accelerates degradation. Overall, these results emphasize a complex relationship between mutation-driven structural changes and post-translational regulation in determining CHIP stability across SCA48 variants.

#### SCA48 mutations cause CHIP mislocalization

An important role of CHIP is to regulate the heat shock response by aiding the trimerization and nuclear movement of heat shock factor 1 (HSF1), which activates heat shock response genes [31]. To assess heat shock responses mediated by CHIP and HSF1, we used a dual-luciferase reporter assay with a plasmid expressing luciferase driven by heat shock elements. Overexpressing WT CHIP increased heat-induced HSF1 transcriptional activity (**Figure 6E**), as heat facilitated the nuclear movement of both HSF1 and CHIP (**Figure 6F-I**). Surprisingly, the selected SCA48 mutants boosted the heat shock transcriptional response similarly to WT (**Figure 6E**), which matched increases in heat-induced HSF1 nuclear translocation (**Figure 6G & I**), indicating this process might be independent of HSC(P) interactions. However, heat shock-induced nuclear translocation of CHIP was missing in all mutants (**Figure 6G-H**). Notably, the normal localization of CHIP at 37 °C was changed, with all mutants showing higher nuclear presence compared to WT CHIP, implying misregulated nucleocytoplasmic shuttling or retention as a potential feature of SCA48 (**Figure 6G-H**). Fractionated immunoblotting and immunofluorescence further supported these results: WT CHIP overexpression increased nuclear localization of both CHIP and HSF1 during heat shock (**Figure 6G-I & Supplementary Figure 3**). The G33S mutant reduced CHIP nuclear localization with a slight defect in HSF1, while P228S and L275Dfs16, despite decreased CHIP localization, maintained or modestly increased HSF1 nuclear levels compared to WT (**Figure 6G-I & Supplementary Figure 3**). This rise in HSF1 activity in U-box mutants may point to a gain-of-toxic-function mechanism. Overall, these findings suggest that CHIP-dependent HSF1 nuclear localization does not strictly depend on HSC(P) interaction but relies on CHIP’s own nuclear import.

#### CHIP expression is necessary for heat-dependent transcriptional responses independent of HSF1 target genes

To further explore CHIP-dependent changes in HSF1-mediated transcriptional activation, we performed RNA-Seq on HEK-293 cells transduced with lentiviruses expressing shRNAs (shControl and sh*STUB1*) under both normal and heat-shock conditions (**Figure 7A**). We observed transcriptional changes in 886 genes in shControl cells and 423 genes in sh*STUB1* cells after heat shock, with 313 genes overlapping. These results suggest significant overlap in heat-stress responses even with CHIP knockdown. Next, we identified HSF1 transcriptional targets from the 313 DEG overlap, revealing remarkably conserved responses between shControl and sh*STUB1* cells after heat shock, with all HSF1-driven changes maintained (**Figure 7B**).

**Figure 7.**
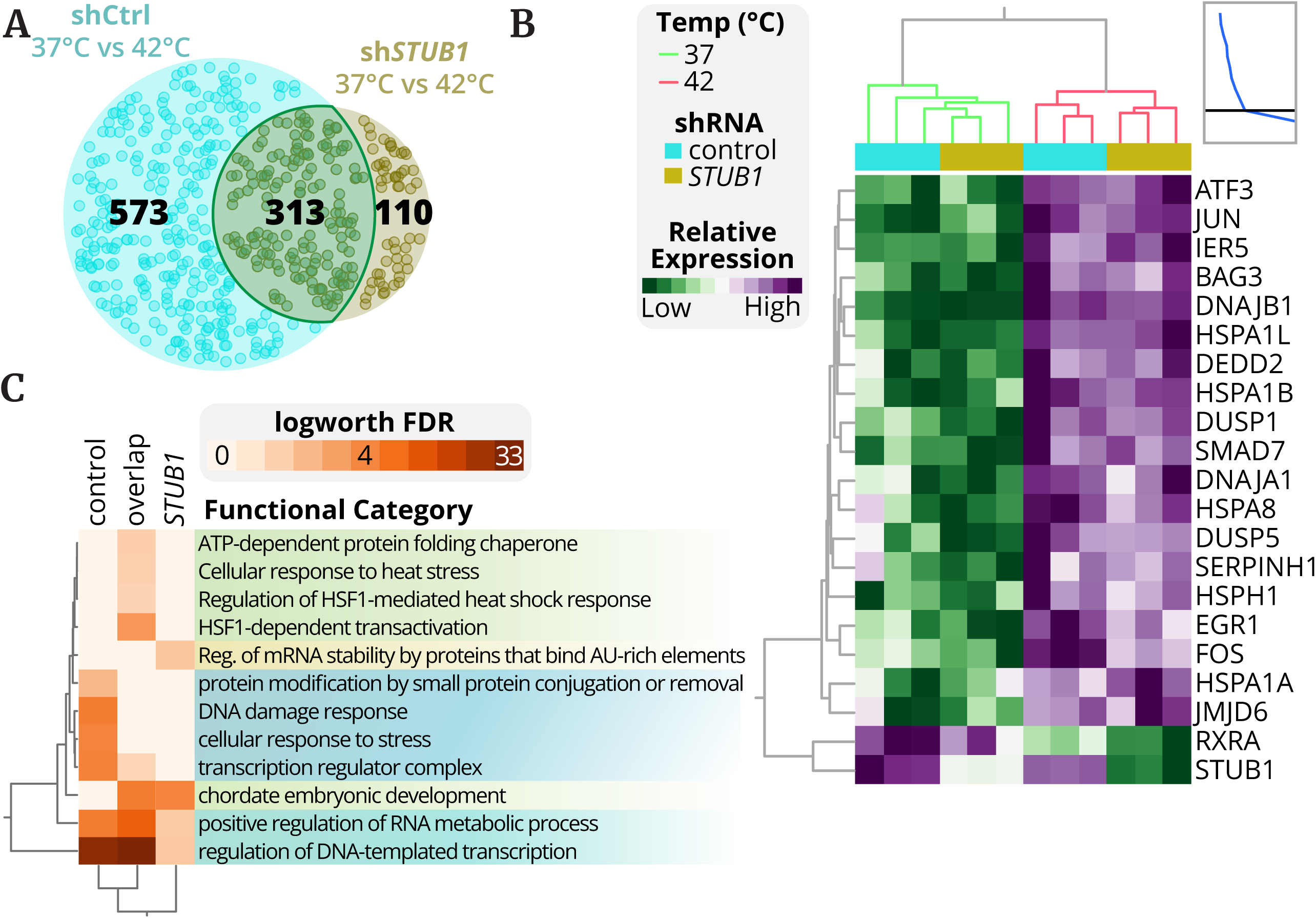
CHIP modulates HSF1-driven transcriptional responses and stress-induced gene networks. (**A**) RNA-sequencing was performed on shCtrl or sh*STUB1* HEK-293 cells under basal conditions and following heat shock. Heat stress-induced differential gene expression (DGE, defined as corrected P < 0.05) identified 886 and 423 genes in shControl and sh*STUB1* cells, respectively, with 313 genes overlapping, summarized by a proportional Venn diagram. (**B**) A heatmap of known critical HSF1 target genes (within the 313-gene overlap) showed largely preserved HSF1-mediated transcriptional activation, including one known heat-stress-suppressed target (RXRA), and confirmed *STUB1* knockdown in the RNAseq data. (**C**) Selected functional enrichment categories of the DEGs from the three Venn diagram regions (in **A**), represented by a hierarchical clustered heatmap of the logworth false discovery rate (FDR).

Functional enrichment analysis of the DEGs further revealed that the overlapping DEG set was enriched for HSF-mediated transcription (**Figure 7C**). Nonetheless, shControl cells exhibited five times more transcriptional changes than sh*STUB1*, with unique enrichments in ATP-dependent protein folding chaperone activity, protein modification by small conjugation or removal, regulation of DNA-templated transcription, positive regulation of RNA metabolic processes, and cellular stress responses, including DNA damage repair. The loss of these categories in sh*STUB1* cells suggests reduced chaperone and ubiquitination activities, consistent with proteostatic defects observed in SCA48, and implies that CHIP enhances the transcriptional heat shock response through mechanisms beyond HSF1, potentially involving nuclear nodes where CHIP modulates DNA damage repair and broader stress pathways.

### Domain-specific correlations with clinical phenotypes in SCA48 patients

Building on the domain-specific biochemical and structural differences observed in SCA48 mutations, we hypothesized that these molecular signatures might be associated with clinical phenotypes. To test this, we conducted a meta-analysis of 87 SCA48 patients from published case studies, gathering data on demographics (age of onset [AOO], sex), mutation locations, Scale for the Assessment and Rating of Ataxia (SARA) scores, and five categorical clinical features representing core symptoms (**Supplementary Table 2**) [4,22,32–36]. The analysis revealed a median AOO of 48 years, with a female majority (about two-thirds of patients; **Figure 8A-B**). Mutations mainly occurred in exons 1-2 (which encode the TPR domain) and 6-7 (which encode the U-box domain), while only 8% were found in exons encoding the coiled-coil (CC) domain (**Figure 8C**). To determine if this distribution was random, we calculated expected mutation frequencies weighted by exon length. Mutations were more frequent than expected in exons 1 (TPR, observed/expected ratio = 1.6, unadjusted/Holm-adjusted p = 0.0166/0.0673) and 7 (U-box, ratio = 1.8, p = 0.0116/0.0579), and less common in exons 3 and 4 (CC, ratio = 0.35 and 0.13; p = 0.0048/0.0335 and 0.0063/0.0376, **Figure 8D**). These differences, tested by chi-square goodness-of-fit while controlling for exon length, suggest exon-specific intolerance: TPR and U-box exons may harbor mutations with higher pathogenicity or be influenced by detection bias, while CC exons face strong selective pressure, possibly due to embryonic lethality or non-viable phenotypes in homozygous models like *Stub1* knockout mice [3]. Compared to SCAR16 (recessive CHIP mutations), where CC domain variants are more common and linked to cognitive deficits and hypogonadism [37], the rarity of CC variants in SCA 48 underscores a dominant-negative mechanism. In this scenario, CC disruptions may completely hinder CHIP dimerization, leading to non-viable or distinct phenotypes. Limitations include potential publication bias favoring symptomatic mutations and limited sequencing depth in CC regions across studies, highlighting the need for larger, uniformly analyzed cohorts.

**Figure 8.**
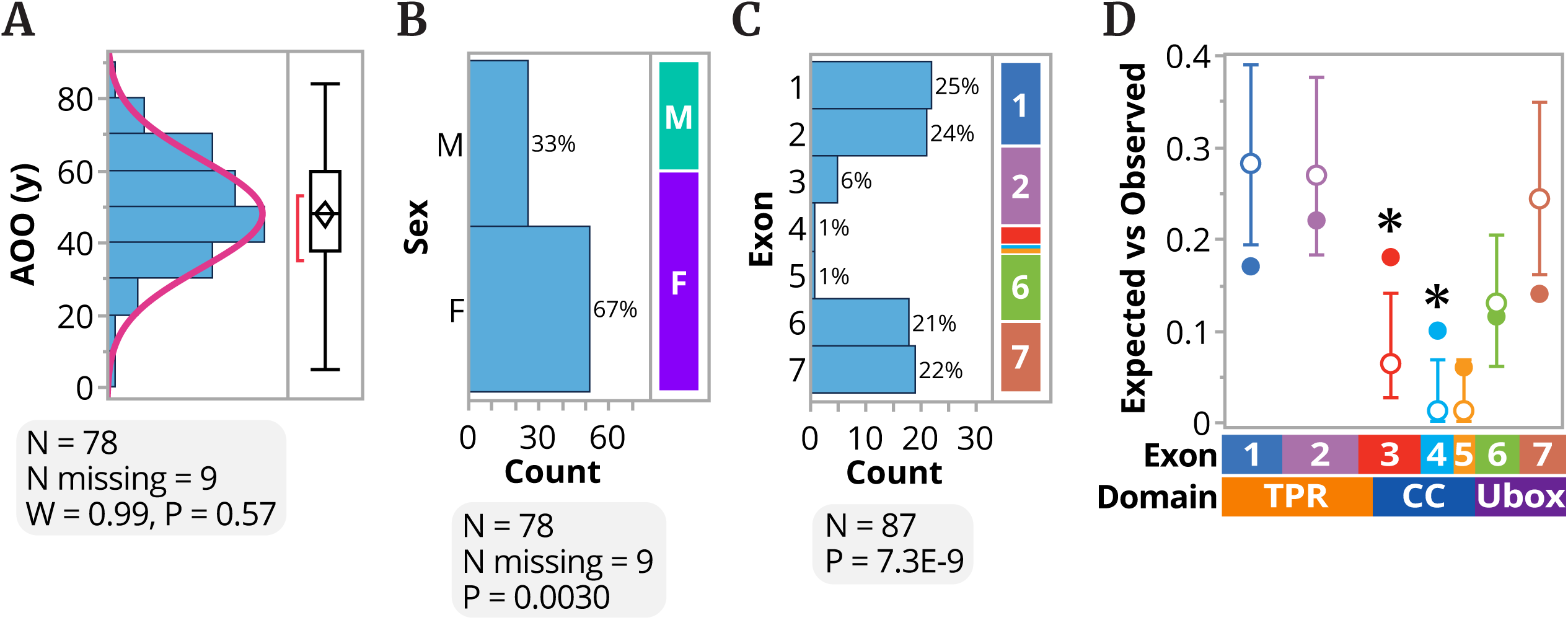
Domain-specific distribution of SCA48 mutations and demographic patterns. (**A**) Distribution (normal, Shapiro-Wilk (W)) of the reported age of onset (AOO) with a median of 48 years, represented by the histogram and box-whisker plot. (**B**) Chi-squared analysis of sex, represented by a contingency plot. (**C**) Distribution of mutations per exon, represented by a contingency plot. (**D**) Mutation mapping across exons represented by plotting the expected (●, ± 95% CI) vs. observed (○) frequency, * P < 0.05.

We then analyzed the relationships between mutation location and clinical phenotypes. U-box mutations were linked to a later age of onset (AOO), with a median of 51 years compared to 42 years for TPR mutations (p = 0.0321; **Figure 9A**), possibly indicating milder early proteostatic effects. However, there was no significant difference in SARA scores between TPR and U-box groups (median 12 vs. 13, p = 0.9490, N = 27; **Supplementary Figure 4**). This suggests that overall disease severity is not solely determined by mutation domain, although factors such as sample size, reporting bias, and variability in assessment timing also influence results. Contingency analysis of categorical symptoms, adjusted for multiple comparisons using false discovery rate (FDR), showed no domain-specific differences in cognitive dysfunction or nystagmus (FDR-adjusted p = 1.0 or 0.6356, respectively; **Supplementary Table 2**). Nevertheless, ataxia was present in 100% of TPR mutations (vs. 86% in U-box; FDR-adjusted p = 0.0344; **Figure 9C**). Dysarthria exhibited a skewed distribution favoring U-box mutations (88% vs. 56% in TPR; FDR-adjusted p = 0.0117; **Figure 9C**). Conversely, TPR mutations were associated with upper motor neuron dysfunction, including increased tendon reflexes (ITR; 78% vs. 25%, FDR-adjusted p = 0.0090; **Figure 9B**) and signs of upper motor neuron involvement (51% vs. 21%, FDR-adjusted p = 0.0344; **Figure 9B**). These FDR-corrected associations (q < 0.05) support the idea that mutation location influences the heterogeneity of the SCA48 spectrum, with TPR mutations predisposing to motor neuron deficits and U-box mutations tending toward dysarthria-dominant presentations. However, limitations include small sample sizes for some features (e.g., ITR N=42), potential variability in clinical reporting across studies, and ascertainment bias toward more severe cases, which restricts broader applicability. Future research with larger, prospective cohorts and standardized phenotyping is needed to validate these findings.

**Figure 9.**
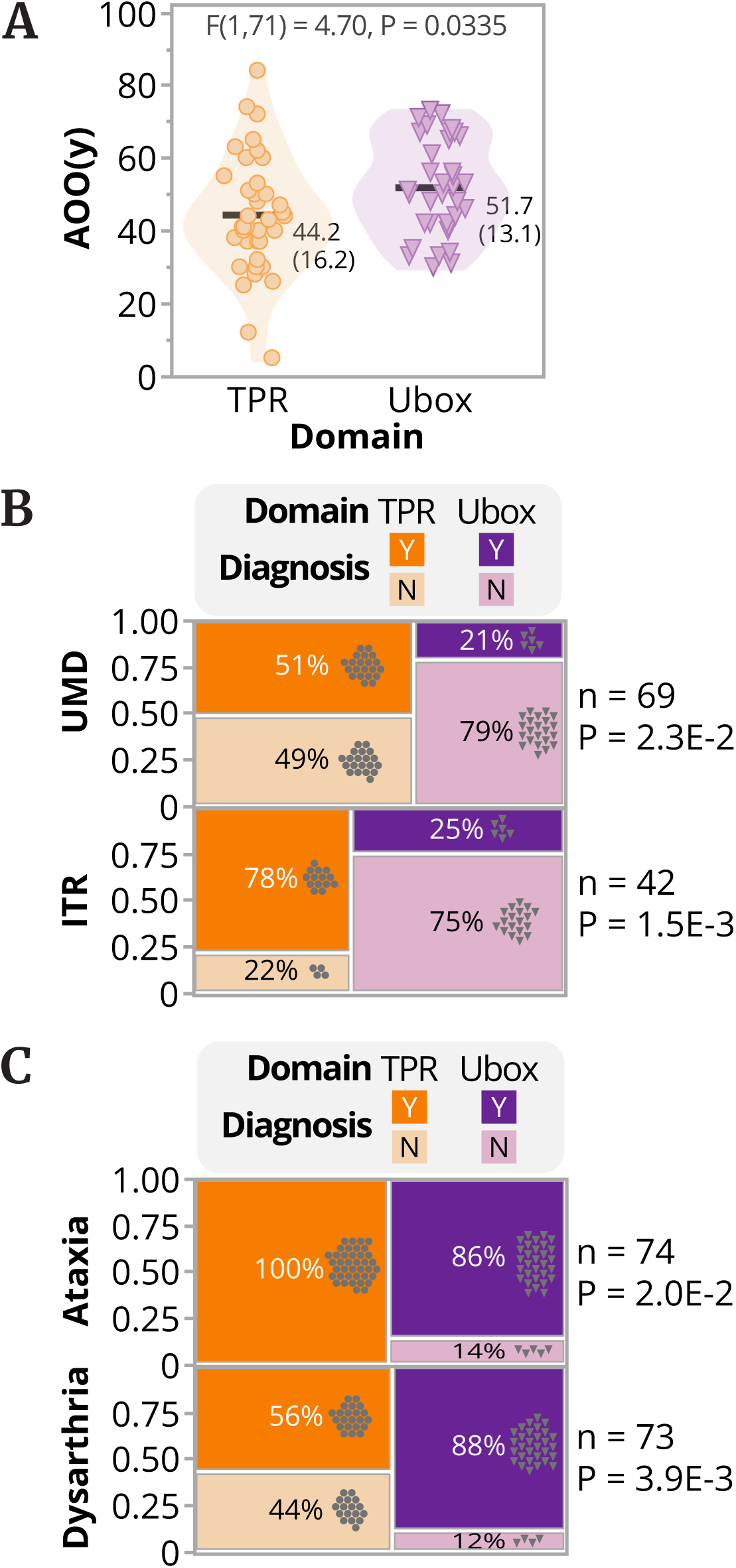
Genotype-phenotype correlations reveal domain-dependent clinical features in SCA48. (**A**) Age of onset (AOO) versus the domain harboring the mutation represented by a violin dot plot, summarized by the mean and standard deviation, analyzed via a two-tailed t-test. (**B & C**) Chi-squared analysis of upper motor-neuron dysfunction (UMD), increased tendon reflexes (ITR), gait ataxia, and dysarthria, represented by a contingency plot. The raw P values of two-tailed Fisher’s exact test are reported, and all fall under the 5% FDR correction for multiple tests.

## Discussion

Our study provides a detailed biochemical and structural framework for understanding how SCA48-associated *STUB1* mutations disrupt CHIP function at multiple levels, including protein stability (**Figures 1 & 6**), post-translational regulation (**Figure 6**), oligomeric assembly (**Figure 2**), and stress response dynamics (**Figures 4, 6, & 7**). These combined findings enhance the mechanistic understanding of how domain-specific mutations interfere with proteostasis pathways and contribute to cellular degeneration. The results emphasize the domain-specific effects of SCA48-related mutations in *STUB1*/CHIP, revealing different mechanisms of dysfunction in the TPR and U-box domains (**Figure 5**). TPR mutations, such as G33S, F37L, I53T, P57L, and A67T, mainly impair substrate recruitment and co-chaperone activity while maintaining the essential ubiquitin ligase function, as shown by decreased HSC70 ubiquitination and luciferase refolding efficiency (**Figures 3 & 4**). Conversely, U-box mutations, including frameshifts and nonsense mutations such as R225*, P228S, and L275Dfs*16, eliminate ligase activity but often preserve or enhance co-chaperone interactions, resulting in increased protein levels and altered oligomerization (**Figures 1, 2, 3, & 4**). These observations align with previous reports on *STUB1* mutations in related ataxias, such as SCAR16, in which U-box disruptions similarly impair ubiquitination without completely disrupting chaperone binding [19,21]. The temperature sensitivity observed in variants such as I53T, A67T, and R241W further suggests that environmental stressors, similar to those that induce mRNA CHIP expression in cells [38], could worsen functional deficits in vivo and may contribute to the progression of neurodegeneration in SCA48 [39]. On the other hand, although ubiquitination defects generally stabilize CHIP by reducing autoubiquitination and proteasomal degradation [22,23], our CHX chase experiments showed that this is not always the case (**Figure 6**). The loss of co-chaperone activity and substrate interaction, often seen in TPR mutants like G33S, resulted in a significantly shorter half-life, suggesting that other factors, such as protein misfolding or changes in structural dynamics, may accelerate turnover. The shorter half-life of G33S and the altered phosphorylation at S19 in P228S U-box mutants suggest post-translational modifications, such as those mediated by protein kinase G [30], may help maintain stability. Consistent with this, variants with lower thermal stability (e.g., G33S, I53T, A67T, and R241W) also exhibit predicted steric clashes in the CHIP dimer interface (**Supplementary Figure 1**), which likely destabilize the overall protein structure and decrease steady-state levels [21,24]. These findings imply that destabilizing mutations disrupt the balance between CHIP formation and degradation, leading to variable protein levels across SCA48 variants.

Biophysical analyses, including differential scanning fluorimetry (**Table 1**) and mass photometry (**Figure 2**), show that SCA48 mutations cause various changes in protein stability and oligomerization, which likely explain the observed functional differences (**Figure 5**). TPR mutants typically form distinct dimer-repeating oligomers (MP Profile 1), allowing some residual activity, whereas most U-box mutants tend to aggregate into non-discrete high-molecular-weight species (MP Profile 3), which correlates with reduced heat-induced oligomer shifts in cellular assays. This tendency to aggregate reflects studies on CHIP’s role in protein quality control, where oligomerization influences chaperone-dependent ubiquitination and refolding [40,41]. Dimerization is critical for co-chaperone binding and ubiquitination, while higher-order oligomers are needed for strong chaperone activity during cellular stress [17]. Mass photometry and biochemical profiling indicate that TPR-domain mutations mostly preserve dimer formation, while U-box variants often produce non-discrete higher-order species or aggregates (**Figure 2**). This change in oligomerization is not just structural; it likely causes downstream effects such as impaired substrate processing and abnormal stress responses. Notably, mutant CHIP’s failure to form higher-order oligomers during heat shock was linked to problems in nuclear translocation and altered heat shock response, highlighting the importance of oligomeric regulation [17,19]. These biochemical defects impact the regulation of HSF1, a key player in the cellular stress response. CHIP promotes HSF1 trimerization and nuclear translocation, boosting the transcription of protective heat shock genes [31,42]. Although SCA48 mutants maintained HSF1 activity under stress, they couldn’t transfer CHIP to the nucleus and, interestingly, showed increased basal nuclear localization (**Figure 6**). This misregulation suggests that CHIP’s movement between the nucleus and cytoplasm is sensitive to structural changes, and abnormal nuclear retention or mislocalization might be a hallmark of SCA48 pathology. Additionally, some U-box mutants, such as L275Dfs*16, still enhance nuclear accumulation of HSF1, suggesting potential gain-of-function effects that could dysregulate the stress response and promote disease progression.

Clinical correlations from the meta-analysis of 87 SCA48 patients show that mutation location affects phenotypic presentation: U-box mutations are associated with later age of onset, and TPR mutations are associated with upper motor neuron signs, including increased tendon reflexes (**Figure 9**). Biochemical profiles, such as low expression, increased autoubiquitination, and decreased co-chaperone activity in TPR variants, support these skewed patterns (**Figure 5**), suggesting that proteostatic imbalances contribute to symptoms such as ataxia and dysarthria (**Figure 9**). This domain-specific pattern resembles genetic modifiers observed in other SCAs, in which *STUB1* variants interact with repeat expansions in genes such as *ATXN8OS*/*ATXN8* or *TBP* to influence penetrance and severity [5,43]. The concentration of mutations in exons encoding TPR and U-box domains, which differs from a random distribution (**Figure 8**), indicates selective pressure on functional hotspots, consistent with CHIP’s dual role in proteostasis and neurodegeneration models [44]. While our data offers valuable insights into CHIP domain-specific functions and their impact on SCA48 clinical features, further research is necessary to understand how other genetic factors, such as intermediate TBP41-49, operate within the SCA48 context and influence the disease phenotype. A recent study of patients with spinocerebellar ataxia 17 (SCA17) and intermediate glutamine repeat expansions (41-49 repeats) in the TATA-box-binding protein (TBP) gene (TBP41-49) identified heterozygous mutations in *STUB1* [6]. Interestingly, while *TBP* alleles with over 49 glutamine repeats are fully penetrant and considered pathological, the TBP41-49 allele shows incomplete penetrance; its frequency in the general population is about 2%, and approximately 50% of carriers of intermediate alleles remain healthy [6]. This raised questions about whether *STUB1* mutations are causal or merely disease-modifying. A follow-up study found normal *TBP* alleles (<40 repeats) in several families with different *STUB1* mutations presenting with the SCA48 phenotype [45]. They also noted an association between dementia and shorter lifespan in patients carrying both *STUB1* mutations and intermediate TBP41-49 alleles, concluding that *STUB1* mutations are causal, while intermediate TBP41-49 alleles act as disease modifiers that lead to a more severe disease presentation [45]. Our clinical association studies excluded patients with mutations in other known ataxia-related genes to better depict the pure *STUB1*-related phenotypes. Although our data provide key insights into CHIP domain-specific functions and their influence on SCA48 clinical presentation, additional research is needed to fully understand how other genetic factors, such as intermediate TBP41-49, operate within the SCA48 framework and contribute to the disease.

The scarcity of mutations in the coiled-coil domain (∼3% of reported SCA48 variants, despite comprising about 15% of *STUB1* length) is notable and suggests strong selective pressure (**Figure 8**). This region, crucial for dimerization [25,46,47], may be highly intolerant to alterations, as disruptions could completely abolish CHIP function and potentially cause embryonic death or non-viable phenotypes, similar to homozygous *Stub1* knockout mice that die perinatally [31,48]. Alternatively, variants in the coiled-coil region might be tolerated but not cause disease in SCA48, thus escaping detection in disease cohorts. Without biochemical or clinical data on these mutations, definitive conclusions are challenging; however, the importance of this domain indicates that targeted sequencing in larger cohorts is necessary to determine whether these mutations are tolerated or lethal.

This domain-clinical mapping reflects patterns observed in other inherited ataxias and motor neuron diseases, where the mutation location influences phenotypic variation [49]. In SCA3 (*ATXN3*), mutations in the N-terminal Josephin domain lead to earlier onset and more severe ataxia, while C-terminal variants near the polyQ tract result in milder forms with notable spasticity or dystonia [50]. Likewise, in SCA2 (*ATXN2*), N-terminal expansions are associated with pure ataxia and slow saccades, whereas rare C-terminal disruptions are linked to Parkinsonism or ALS-like features. In ALS (*SOD1*), mutations in the N-terminal metal-binding domain often cause rapid progression with lower motor neuron (LMN) dominance, while C-terminal variants tend to have slower courses with upper motor neuron (UMN) signs [51–54]. In hereditary spastic paraplegia type 4 (SPG4), *SPAST* missense mutations in the C-terminal ATPase (AAA) domain are typically associated with pure spastic paraplegia primarily affecting upper motor neurons, whereas truncating or frameshift variants, often disrupting the overall protein function, including the microtubule-binding domain, are linked to complex phenotypes with additional features such as ataxia and dysarthria [55–59]. These parallels support SCA48’s model: TPR (upstream recognition) leads to aggressive UMN/ataxia symptoms, similar to N-terminal disruptions, while U-box (downstream ligase) results in more subtle bulbar involvement, resembling C-terminal effects.

Pathophysiologically, the universal cerebellar atrophy in SCA48 indicates a common core mechanism [32,39]: dominant mutants poison wild-type dimers, impair proteostasis, and cause aggregate buildup in Purkinje and granule cells, regardless of domain. Divergence occurs from domain-specific differences: for example, TPR mutations may trap substrates, causing ER stress and UPR in UMN pathways (explaining high reflexes, dysfunction, and early onset), while U-box failures lead to chaperone hyperactivity, which buffers ataxia but strains bulbar neurons (resulting in high dysarthria and ataxia-sparing) [60,61]. Regarding UMN versus bulbar involvement, motor neuron diseases like ALS show proteostasis-related susceptibility: UMN vulnerability often stems from cortical ER stress due to trapped aggregates, consistent with TPR’s client bottleneck, while bulbar/LMN strain involves chaperone overload and axonal transport issues, matching CHIP’s U-box failure [62,63]. The lack of difference in SARA severity (despite a small sample size of N=27) supports cerebellar convergence, but larger groups might reveal subtle gradients [64,65].

Finally, these insights open the door for targeted therapies in SCA48 and related ubiquitin ligase disorders [66]. Stabilizing TPR-client interactions could lessen early proteotoxic stress, similar to how ATXN3 inhibitors that target deubiquitinase activity reduce aggregate formation and improve motor symptoms in SCA3 models [67,68]. For U-box mutations, restoring ligase activity with small-molecule activators of E3 ligases (like PROTACs that recruit CHIP) or chaperone modulators (such as HSP70 inhibitors to reduce overload) shows promise [69,70]. This approach has been demonstrated in ALS trials that target SOD1 ubiquitination and in Alzheimer’s disease models where PROTACs degrade tau aggregates by recruiting E3 ligases like VHL or CRBN, thereby decreasing neurotoxicity [71,72]. Broader drugs that target the ubiquitin pathway, for example UCHL1 inhibitors for neurodegeneration, could link different domains. In Parkinson’s disease models, UCHL1 inhibition lowers alpha-synuclein aggregation, and in Alzheimer’s, it reduces tau pathology by boosting lysosomal clearance [73,74]. Future therapies might use CRISPR-based domain editing or antisense oligonucleotides (ASOs) to target mutant alleles. For instance, CRISPR-Cas9 suppression of mutant ATXN3 in SCA3 models improves neuronal deficits [75], and ASO-mediated silencing in Friedreich’s ataxia restores frataxin levels and enhances sensory functions [76], potentially transforming SCA48 from an untreatable disorder into a condition that can be managed therapeutically.

## Materials and Methods

### Plasmids, Mutagenesis, and Protein Purification

We used the pcDNA3-CHIP-myc and pET30-CHIP-His plasmids for mammalian and bacterial expression, respectively, as described earlier [19]. Disease-related mutations were introduced using the Q5® Site-Directed Mutagenesis Kit (New England Biolabs, E0554S) following the manufacturer’s instructions, with primers listed in **Supplementary Table 3**.

CHIP mutants were expressed with a C-terminal His tag in BL21 (DE3)-RIL cells. Cells were resuspended and lysed in 1X PBS, 20 mM imidazole, 5 mM BME. Proteins for mass photometry were purified via nickel affinity chromatography and buffer exchanged into 20 mM HEPES pH 8.0, 200 mM NaCl, 1 mM DTT using Cytiva PD-10 desalting columns. Proteins for ubiquitination assays were purified via nickel affinity chromatography followed by size exclusion chromatography in 20 mM HEPES pH 8.0, 200 mM NaCl, 1 mM DTT.

Fluorescently labeled HSC70 (*HSC70) was prepared using a GST-tagged, sortase-compatible Hsc70 construct. This construct was expressed similarly to CHIP but was resuspended and lysed in 50 mM Tris pH 7.6, 200 mM NaCl, and 5 mM BME. The protein was purified through glutathione affinity chromatography, and the GST tag was cleaved with PreScission-GST. Uncleaved protein was removed by passing it through glutathione affinity chromatography again. GST-cleaved HSC70, which contains an N-terminal Glycine residue, was fluorescently labeled via an overnight reaction with 5 µM protein, 50 µM fluorescent peptide, and 5 µM His-sortase. The reaction was quenched with 10 mM EDTA, then passed through a Cytiva PD-10 desalting column and nickel affinity chromatography to remove excess peptide and His-sortase. Finally, *HSC70 was further purified by size exclusion chromatography using 20 mM HEPES pH 8.0, 200 mM NaCl, and 1 mM DTT.

### Cell Culture, Transfection, and Heat Shock Induction

COS-7 and HEK293 cells were cultured in Dulbecco’s Modified Eagle’s Medium (Corning, 10-017-CV) supplemented with 10% fetal bovine serum (Gibco, 26140-079) and incubated at 37°C in a 5% CO2 atmosphere. Transient transfections were performed using X-tremeGENE™ 9 DNA transfection reagent (Roche, 6365787001) following the manufacturer’s instructions, with a 1:3 DNA to reagent ratio. Heat shock was applied by incubating cells at 42 °C for two hours in a 5% CO2 environment. Stable knockdown of STUB1/CHIP was achieved using a combination of four validated shRNA lentiviruses (TRCN0000007525, TRCN0000007526, TRCN0000007528, TRCN0000007529).

### Mass Photometry

Mass photometry was conducted using a Refeyn OneMP Mass Photometer placed on an Accurion i4 vibration isolation system. Samples were prepared by washing ThorLabs Precision microscope cover slips through sonication for 10 minutes in a 50:50 mixture of Isopropanol and DI water, followed by 10 minutes in 100% DI water. The cleaned cover slips were stored in DI water and dried immediately before use with filtered air. CultureWell™ gaskets (Grace Bio-Labs) were glued to the dried cover slips. The mass photometer lens was prepped by adding a drop of Olympus IMMOIL-F30CC immersion oil, after which the prepared cover slip was placed on top. For each well, 15 µL of filtered 20 mM HEPES pH 8.0 with 200 mM NaCl buffer was added, and the lens was focused near the edge of the well using the droplet-dilution method. Once proper focus was achieved, 5 µL of a 200 nM prediluted CHIP sample was added to the well, resulting in a total volume of 20 µL and a final concentration of 50 nM. The signal was briefly allowed to stabilize, then a 60-second video was recorded at 100 frames per second using the AcquireMP software. The video was processed in DiscoverMP, and a mass calibration curve was created using BSA (Sigma P0834) and ApoFerritin (Sigma A3660) to estimate the mass of each event. The analyzed events were then processed in Python to generate normalized bar and scatter plots, which were visualized using GraphPad Prism.

### HSC70 Ubiquitination and Autoubiquitination Assays

HSC70 ubiquitination assays were performed using HSC70 tagged with a fluorescent N-terminus. All assay components except CHIP were premixed on ice in a BSA-containing buffer (20 mM HEPES, pH 8.0, 200 mM NaCl, 0.5 mg/mL BSA). Reactions were initiated by adding CHIP and kept at room temperature unless otherwise specified. The reactions contained 1 µM Uba1, 25 µM UBE2D2, 1 µM *HSC70, 100 µM Ub, 5 mM MgCl2-ATP, and 5 µM CHIP. They were allowed to proceed for 15 minutes before being quenched with SDS loading buffer. The quenched samples were separated by molecular weight using SDS-PAGE under non-reducing conditions with GenScript SurePAGE 4-12% gels. Fluorescent scanning on an Amersham Typhoon 5 enabled visualization and quantification of CHIP-dependent ubiquitination of *Hsc70.

CHIP autoubiquitination was monitored through pulse-chase assays. First, a 6x pulse mixture of 0.1 µM Uba1, 10 µM UBE2D2, 10 µM *Ub, and 5 mM MgCl2-ATP was mixed on ice to form UBE2D2∼*Ub (“∼” denotes a thioester intermediate). The UBE2D2∼*Ub mixture was incubated at room temperature for 10 minutes, then quenched with 50 mM EDTA. Chase mixes were prepared with 5 µM of either wild-type or variant CHIP, and the reaction was started by adding the pulse mix to a 1x concentration. The reactions were allowed to proceed at room temperature and quenched with SDS loading buffer at the indicated time points. Quenched samples were analyzed by SDS-PAGE and fluorescent scanning, as described above for substrate ubiquitination assays.

### Cell Lysis and Soluble-Insoluble Fractionation

Cells were maintained and transfected as described above. Twenty-four hours after transfection, cells were lysed in ice-cold 1% Triton X-100 buffer (50 mM Tris-HCl pH 7.4, 150 mM NaCl, 2 mM EDTA, 1% Triton X-100) supplemented with 1X Halt protease and phosphatase inhibitor (Thermo Scientific, 78447). Lysates were cleared by centrifugation at 15,000 rpm for 10 minutes. BCA assay (Thermo Scientific, 23223 & 23224) was performed to determine protein concentrations, and 50 µg of total protein was used for immunoblotting.

For soluble and insoluble fractionation, cells were lysed in ice-cold 1% Triton X-100 buffer (50 mM Tris-HCl pH 7.4, 150 mM NaCl, 2 mM EDTA, 1% Triton X-100) supplemented with 1X Halt protease and phosphatase inhibitors (Thermo Scientific, 78447). Lysates were cleared by centrifugation at 16,000 x g for 30 minutes at 4 °C. The supernatants (Triton X-100 soluble fraction) were transferred to a fresh tube. The pellets were washed with lysis buffer and centrifuged at 16,000 x g for 10 minutes at 4 °C. Lysis buffer was discarded, and the pellets were resuspended in 1X Laemmli sample buffer (Bio-Rad, 1610747). Samples were briefly sonicated, and protein concentrations were measured (Thermo Scientific, 22660). Fifty micrograms of Triton X-100 soluble and insoluble fractions were used for immunoblotting.

### Immunoblotting, Immunoprecipitation, and Antibodies

Cells were lysed as described above, and protein concentrations were measured using the BCA Protein Assay Kit (Thermo Scientific, 23223 & 23224). Equal amounts of total protein (15-50 µg) were mixed with 1X Laemmli sampling buffer (Bio-Rad, 1610747) and boiled at 100°C for 5 minutes. Proteins were separated on 4-15% Mini-PROTEAN TGX Stain-Free gels (Bio-Rad, 4568084) and then transferred to PVDF membranes (Bio-Rad, 1620264). Membranes were blocked with 5% milk or bovine serum albumin (BSA) and incubated overnight at 4°C with primary antibodies diluted in the appropriate blocking solution (**Supplementary Table 4**). After washing in TBST, membranes were incubated with HRP-conjugated anti-rabbit secondary antibody (Cell Signaling Technology, 7074), and visualized using Lumigen ECL Ultra (Lumigen, tma-100). For fluorescent western blot detection, primary antibodies against phospho-CHIP (Ser20) (Abmart Inc., 25011-1) and MYC (Proteintech, 60003-2-Ig) were used, followed by fluorescent secondary antibodies StarBright 700 (Bio-Rad, 12004161) or StarBright 520 (Bio-Rad, 12005866). For immunoprecipitation, 10E6 cells were plated in 10 cm tissue culture dishes and transfected as described above. Twenty-four hours post-transfection, cells were lysed in ice-cold 1% Triton X-100 buffer (50 mM Tris-HCl pH 7.4, 150 mM NaCl, 2 mM EDTA, 1% Triton X-100) supplemented with 1X Halt Protease and Phosphatase Inhibitor Cocktail (Thermo Scientific, 78447). Lysates were cleared by centrifugation at 15,000 rpm for 10 minutes, and protein concentrations were measured using a BCA assay. For each sample, 1 mg of total protein was used for immunoprecipitation. CHIP-myc and Flag-HSP70 were pulled down with Ezview Red Anti-c-Myc affinity gel (Sigma-Aldrich, E6654) and Ezview Red Anti-Flag M2 affinity gel (Sigma-Aldrich, F2426), respectively, following the manufacturer’s instructions. Cell lysates were incubated with affinity gels overnight at 4 °C with rotation. Beads were washed five times before elution with 1X Laemmli sampling buffer in 1% Triton X-100 buffer and boiled at 100 °C for 5 minutes. Eluted samples and inputs were run on 4-15% Mini-PROTEAN TGX Stain-Free gels and transferred to PVDF membranes for immunoblotting as described above.

### Immunofluorescence

COS-7 cells were fixed with 4% paraformaldehyde (PFA) for 15 minutes at room temperature, then washed three times with 1× phosphate-buffered saline (PBS). Fixed cells were either stored at 4 °C or processed immediately for immunofluorescence staining. For blocking and permeabilization, cells were incubated for 60 minutes at room temperature in 5% bovine serum albumin (BSA; Fisher Scientific, BP9703-100) diluted in PBS containing 0.1% Triton X-100. Subsequently, cells were incubated overnight at 4 °C with primary antibodies against Myc (Proteintech, 60003-2-Ig, D1I4Q) and HSF1 (Cell Signaling Technology, #4356), both diluted in 1% BSA with 0.1% Triton X-100.

Following primary antibody incubation, cells were washed three times with PBS containing 0.1% Triton X-100 for 10–15 minutes each. Samples were then incubated for 1 hour at room temperature with Multi-rAb™ CoraLite® Plus 488-Goat Anti-Rabbit Recombinant Secondary Antibody (H+L) and Multi-rAb™ CoraLite® Plus 555-Goat Anti-Mouse Recombinant Secondary Antibody (H+L) (Proteintech), both diluted in 1% BSA with 0.1% Triton X-100. After secondary antibody incubation, cells were rinsed three times with PBS containing 0.1% Triton X-100. During the second wash, NucBlue™ Fixed Cell ReadyProbes™ Reagent (Invitrogen, MA, USA) was added to stain nuclei.

After the final wash, samples were mounted on glass slides and covered with #1.5 coverslips using ProLong™ Diamond Antifade Mountant (Thermo Fisher Scientific, NC, USA). Immunofluorescence images were captured with a Nikon Eclipse Ti2 microscope equipped with a 40×/0.95 NA Plan Apochromat air objective and controlled by Nikon Elements software. Fluorescence signals were recorded using a pco.edge 4.2Q High QE sCMOS camera, providing high sensitivity and resolution for quantitative image analysis.

#### Image analysis

Images were processed and analyzed using the ImageJ/FIJI software. The nuclear localization of Myc and HSF1 was quantified by comparing nuclear to cytoplasmic intensity ratios or by normalizing nuclear Myc or HSF1 intensity to the total nuclear signal (for example, nuclear Myc/nuclear NucBlue intensity). In the figures, the nuclear intensity of Myc or HSF1 is shown using a masked nuclear region created based on NucBlue™ nuclear staining, with the intensity displayed in a 16-color LUT (FIJI).

### Blue Native Polyacrylamide Gel Electrophoresis

COS-7 cells were maintained and transfected as described above. Twenty-four hours after transfection, samples were collected in Native PAGE sample buffer (Invitrogen, BN2003) supplemented with 1X Halt Protease and Phosphatase Inhibitors cocktail (Thermo Scientific, 78447) and 1% digitonin (Calbiochem, 300410). Lysates were clarified by centrifugation at 20,000 × g for 30 minutes at 4 °C. Protein concentrations were measured using a BCA assay. G-250 Sample Additive (Invitrogen, BN2004) was added to the samples, and blue native PAGE was run using the dark blue and light blue cathode buffer protocol according to the manufacturer’s instructions (Invitrogen). Native protein samples were resolved on 18-well 4-15% Criterion TGX Stain-Free Precast Gels (Bio-Rad, 5678084) using 0.25% G-250 cathode buffer (Invitrogen, BN2007) on ice and transferred to PVDF membrane. Immunoblot analysis was performed as described above.

### Cycloheximide Chase Assay

COS-7 cells were maintained as described above. Twenty-four hours post-transfection, cells were treated with 50 µg/ml cycloheximide at the specified time points and then harvested using Triton X-100 buffer. Fifty µg of total protein were separated by SDS-PAGE as described earlier, followed by immunoblot analysis.

### Luciferase Refolding Assay

Recombinant CHIP, HSC70, and HSP40 proteins were purified as described above. Recombinant luciferase (Promega, E1701) was prepared at 0.2 µM in 2 mg/ml BSA (Thermo Fisher Scientific, BP9706100) and heat-denatured by incubation at 42 °C for 15 minutes. Recombinant CHIP, HSC70, and HSP40 proteins were diluted to 32 µM in HEPES protein storage buffer. Denatured luciferase and recombinant proteins were diluted 1:4 in protein storage buffer and incubated at 37°C for an hour to allow refolding. Three replicates of 2 µl reaction mix were aliquoted into a white opaque 96-well plate. A CLARIOstar plate reader (BMG Labtech) was used to inject 50 µl Luciferase Assay Reagent (Promega, E1483) into each well, followed by luminescence detection.

### HSF1 Luciferase Reporter Assay

HSF1 luciferase reporter assay was conducted as previously described [19]. Briefly, cells were transfected with the indicated vectors alongside the Qiagen Heat Shock Response Reporter Kit (Qiagen, CCS4023L) according to the manufacturer’s instructions. Twenty-four hours after transfection, cells were lysed with 1X passive lysis buffer (Promega, E1910), and the luciferase assay was performed using the Dual-Luciferase Reporter Assay System (Promega, E1910) on a CLARIOstar plate reader following the manufacturer’s protocol. Three biological replicates, each with three technical replicates, were conducted on separate days. Luciferase/Renilla ratios were calculated, and technical replicates were averaged. Ratios were normalized to wells transfected with empty vector.

### Differential Scanning Fluorometry

Recombinant CHIP proteins were purified as described above. Reaction buffers were prepared by diluting the indicated CHIP proteins to 20 µM in 10X SYPRO Orange (Sigma-Aldrich, S5692) in HEPES protein storage buffer. Three replicates of 10 µl reaction buffer were aliquoted into 384-well PCR plates, and the melting temperature assay was performed on a LightCycler 480 (Roche) following the manufacturer’s protocol.

Fluorescence was measured at 568 nm while the temperature was ramped from 20 °C to 85 °C with 10 acquisitions per degree Celsius. The melting temperatures of CHIP proteins were determined by calculating the first derivative of the fluorescence data and identifying the local maxima.

### Densitometry Analysis

Immunoblot quantification was performed using densitometry analysis with BioRad Image Lab software (v6.1).

### RNA Sequencing and Differential Expression Analysis

#### Cell culture, heat shock, and RNA isolation

HEK-293 cells were transduced with lentiviral particles expressing either a non-targeting control shRNA (shControl) or an shRNA targeting *STUB1* (sh*STUB1*). Stable cell populations were selected with puromycin (2 µg/mL) for 72 h and cultured in DMEM supplemented with 10% FBS and 1% penicillin-streptomycin at 37 °C in 5% CO₂. Heat shock was induced by incubation at 42 °C for 30 minutes, and the cells were immediately collected for RNA purification. Three independent biological replicates were generated for each of four conditions: shControl 37 °C, shControl 42 °C, sh*STUB1* 37 °C, and sh*STUB1* 42 °C (n = 12 samples total). Total RNA was isolated using the RNeasy Mini Kit (Qiagen) with on-column DNase I digestion. RNA purity was assessed using NanoDrop (A260/A280 ≥ 2.0, A260/A230 ≥ 1.8); integrity (RIN ≥ 9.0) was confirmed with the RNA 6000 Nano Kit on a 2100 Bioanalyzer (Agilent).

#### Library construction and sequencing

Poly(A)-selected mRNA libraries were prepared using the Illumina TruSeq RNA Sample Preparation Kit. Sequencing was performed single-end 50 cycles on an Illumina HiSeq 2500, producing 15.8–36.4 million read pairs per sample (average: 24.7 million). Raw data quality was confirmed using FastQC v0.11.9 (per-base Phred score >30 in at least 95% of cycles).

#### Read alignment and quantification

Reads were aligned to the human reference genome (GRCh38.p13) using STAR v2.7.8a in 2-pass mode with default parameters. Alignment was guided by Ensembl transcript annotations (release 105). Gene-level read counts were quantified using Partek Expectation/Maximization (E/M) in Partek Flow v10.0.23.0214, counting only uniquely mapped, properly paired reads overlapping annotated exons. Initial quantification included 27,173 genes.

*Differential expression analysis* was performed in Partek Flow v10.0.23.0214 using the DESeq2 node with median ratio normalization. A low-count filter was applied: genes with less than 1 count per million (CPM) in at least 30% of samples were removed, resulting in 15,475 genes retained (11,698 genes were filtered out). The linear model included genotype (shControl, sh*STUB1*), temperature (37 °C, 42 °C), and their interaction.

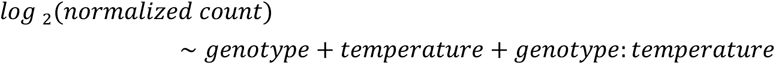

Wald tests were performed for the following contrasts:

- shCHIP vs. shControl (37 °C)
- 42 °C vs. 37 °C (shControl)
- 42 °C vs. 37 °C (shCHIP)
- shCHIP 42 °C vs. shControl 42 °C
- shCHIP 37 °C vs. shControl 37 °C
- Interaction: (shCHIP 42 °C vs. shCHIP 37 °C) vs. (shControl 42 °C vs. shControl 37 °C)

FDR step-up (Benjamini–Hochberg) correction was applied. Independent filtering was enabled; fold change shrinkage was not used. Differentially expressed genes (DEGs) were identified as having FDR < 0.05 in heat shock contrasts (42 °C vs. 37 °C). This process found 886 heat-responsive genes in shControl (573 unique), 423 in sh*STUB1* (110 unique), and 313 common genes (overlap). Variance-stabilized counts were used for downstream visualization.

#### Functional enrichment analysis

Three gene sets were analyzed using g:Profiler (g:GOSt functional profiling, ens109) in multi-query mode:

1. **shControl-unique DEGs** (n = 573)
2. **shCHIP-unique DEGs** (n = 110)
3. **Overlap DEGs** (n = 313)

The background gene set included 15,476 filtered genes. Enrichment analysis was performed using Gene Ontology (Biological Process, Molecular Function, Cellular Component) and Reactome pathways, with pathways identified via Grok AI (xAI) to filter and prioritize results using an FDR threshold of <0.05. To focus on pathways relevant to SCA48, we targeted processes consistent with known pathophysiology, such as protein homeostasis, stress response, ubiquitination, and neuronal dysfunction, as characterized by CHIP (*STUB1*) mutations in SCA48. This involved highlighting categories with significant adjusted p-values and intersection sizes in the overlap (where CHIP function is preserved) and control groups, compared to reduced significance in shCHIP, indicating loss of function. Particular attention was given to HSF1/heat response conservation in the overlap and to unique enrichments that distinguish control from shCHIP conditions. Results passing an FDR < 0.05 cutoff were visualized as a heatmap displaying −log₁₀(FDR) values across the three conditions.

#### Identification of HSF1 target genes

Candidate HSF1 targets were identified by systematically cross-referencing multiple high-confidence, publicly available sources of direct HSF1 targets in human cells. Direct targets were defined as genes with evidence of HSF1 binding (ChIP-seq/ChIP-chip peaks near the transcription start site) and/or HSF1-dependent transcriptional regulation under proteotoxic stress (heat shock, proteostasis challenge) or in cancer contexts. Genes were retained only if supported by at least two independent sources (ENCODE, ChIP-Atlas, HSF1Base, and the non-canonical HSF1 regulon) to ensure robustness and reduce false positives [77–84].

### Meta-analysis of SCA48 clinical phenotypes

#### Data collection and cohort assembly

Clinical and genetic data from 87 SCA48 patients with heterozygous *STUB1* mutations were combined from eight published studies across various geographical locations (**Supplementary Table 2**). Inclusion required a confirmed pathogenic variant (missense, frameshift, nonsense, or splice-site), at least one documented clinical feature, and a diagnosis of cerebellar ataxia. Variables extracted included age of onset (AOO, years), sex, *STUB1* cDNA and protein changes, exon, domain, SARA score, presence of gait ataxia (Y/N), dysarthria (Y/N), cognitive dysfunction (Y/N), upper motor neuron dysfunction (UMND; Y/N), increased tendon reflexes (ITR; Y/N), and nystagmus (Y/N). Cerebellar atrophy was consistently reported (100%) when used as a clinical diagnostic criterion, but was not included in association analyses. Missing data were marked as N/A and excluded from per-feature statistical tests (Table S2.5).

#### Mutation annotation and domain assignment

Variants were annotated relative to the canonical *STUB1* transcript (ENST00000219548.9). Domain boundaries were assigned based on structural and functional data: TPR (aa 1–126), CC (aa 127–221), and U-box (aa 222–303). Exon/AA boundaries are 1 (1–53), 2 (54–120), 2 (121–175), 4 (176–204), 5 (205–223), 6 (224–262), and 7 (163–303). Frameshift and nonsense variants were classified based on the domain containing the premature termination codon.

#### Statistical analyses

Analyses were performed using JMP Pro v19 (SAS Institute). Two-sided p-values less than 0.05 were considered statistically significant.

1. <u>Mutation distribution and exon intolerance.</u> Expected mutation frequencies per exon were calculated as:

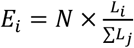 where *N*= total mutations and *L*_*i*_= coding length of exon *i*. Observed vs. expected counts were compared using chi-square goodness-of-fit with Bonferroni correction across seven exons.
2. <u>Age of onset and disease severity.</u> AOO was compared between TPR and U-box mutation groups using a two-tailed Student’s t-test, which was confirmed to be normal by the Shapiro–Wilk test (W = 0.98, p = 0.4921). SARA scores were categorized into five severity levels (minimal: 1–9; moderate: 10–12; maximal: 13–17; severe: 18–24; total: 25–40). Domain associations were analyzed with a contingency table using Fisher’s exact test.
3. <u>Categorical symptom associations.</u> Six binary clinical features (ataxia, dysarthria, cognitive dysfunction, UMND, ITR, nystagmus) were tested against the domain (TPR vs. U-box) using 2×2 contingency tables and Fisher’s exact test. P-values were adjusted with the false discovery rate (FDR; Benjamini–Hochberg) across the tests.
4. <u>Sensitivity and Limitations.</u> Due to the retrospective, multinational design, variation in clinical assessment timing, diagnostic criteria, and reporting depth was inevitable. Results should be viewed as hypothesis-generating, pending validation in prospective, standardized cohorts.

### Statistical Analysis Software

All statistical analyses for the biochemical and clinical data were performed using GraphPad Prism 9 and JMP Pro (v19.0.0), respectively.

## Supplemental Information

### Supplementary Tables

**Supplementary Table 1.**
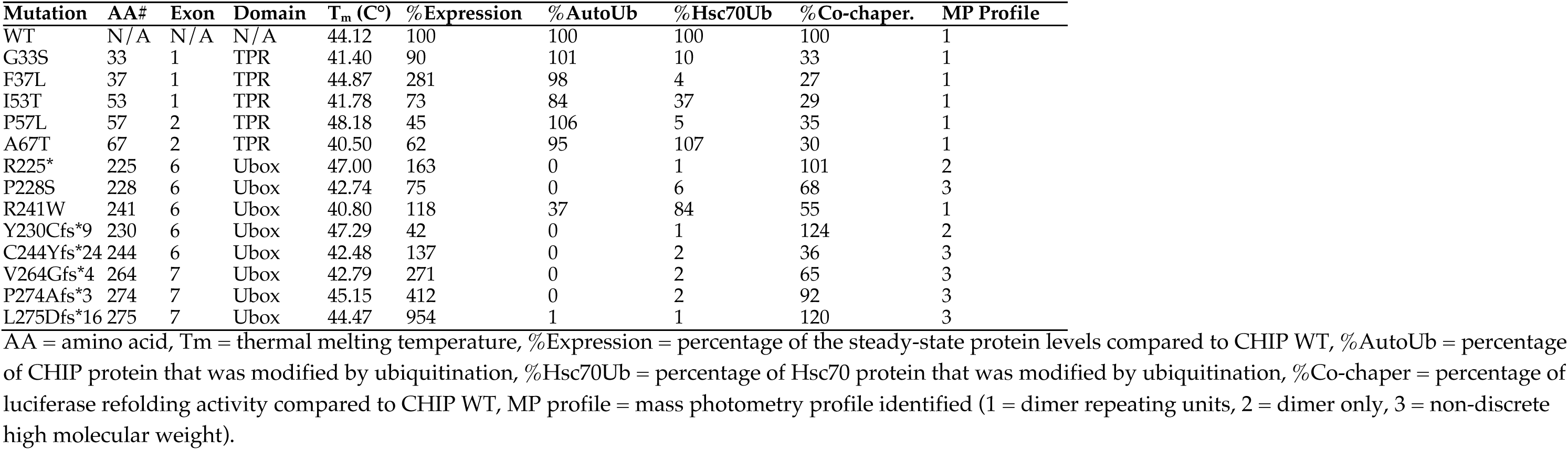
Biochemical Profiles of CHIP mutants.

| Mutation | AA# | Exon | Domain | T<sub>m</sub> (C°) | %Expression | %AutoUb | %Hsc70Ub | %Co-chaper. | MP Profile |
| --- | --- | --- | --- | --- | --- | --- | --- | --- | --- |
| WT | N/A | N/A | N/A | 44.12 | 100 | 100 | 100 | 100 | 1 |
| G33S | 33 | 1 | TPR | 41.40 | 90 | 101 | 10 | 33 | 1 |
| F37L | 37 | 1 | TPR | 44.87 | 281 | 98 | 4 | 27 | 1 |
| I53T | 53 | 1 | TPR | 41.78 | 73 | 84 | 37 | 29 | 1 |
| P57L | 57 | 2 | TPR | 48.18 | 45 | 106 | 5 | 35 | 1 |
| A67T | 67 | 2 | TPR | 40.50 | 62 | 95 | 107 | 30 | 1 |
| R225* | 225 | 6 | Ubox | 47.00 | 163 | 0 | 1 | 101 | 2 |
| P228S | 228 | 6 | Ubox | 42.74 | 75 | 0 | 6 | 68 | 3 |
| R241W | 241 | 6 | Ubox | 40.80 | 118 | 37 | 84 | 55 | 1 |
| Y230Cfs*9 | 230 | 6 | Ubox | 47.29 | 42 | 0 | 1 | 124 | 2 |
| C244Yfs*24 | 244 | 6 | Ubox | 42.48 | 137 | 0 | 2 | 36 | 3 |
| V264Gfs*4 | 264 | 7 | Ubox | 42.79 | 271 | 0 | 2 | 65 | 3 |
| P274Afs*3 | 274 | 7 | Ubox | 45.15 | 412 | 0 | 2 | 92 | 3 |
| L275Dfs*16 | 275 | 7 | Ubox | 44.47 | 954 | 1 | 1 | 120 | 3 |

AA = amino acid, T<sub>m</sub> = thermal melting temperature, %Expression = percentage of the steady-state protein levels compared to CHIP WT, %AutoUb = percentage of CHIP protein that was modified by ubiquitination, %Hsc70Ub = percentage of Hsc70 protein that was modified by ubiquitination, %Co-chaper = percentage of luciferase refolding activity compared to CHIP WT, MP profile = mass photometry profile identified (1 = dimer repeating units, 2 = dimer only, 3 = non-discrete high molecular weight).

**Supplementary Table 2.**
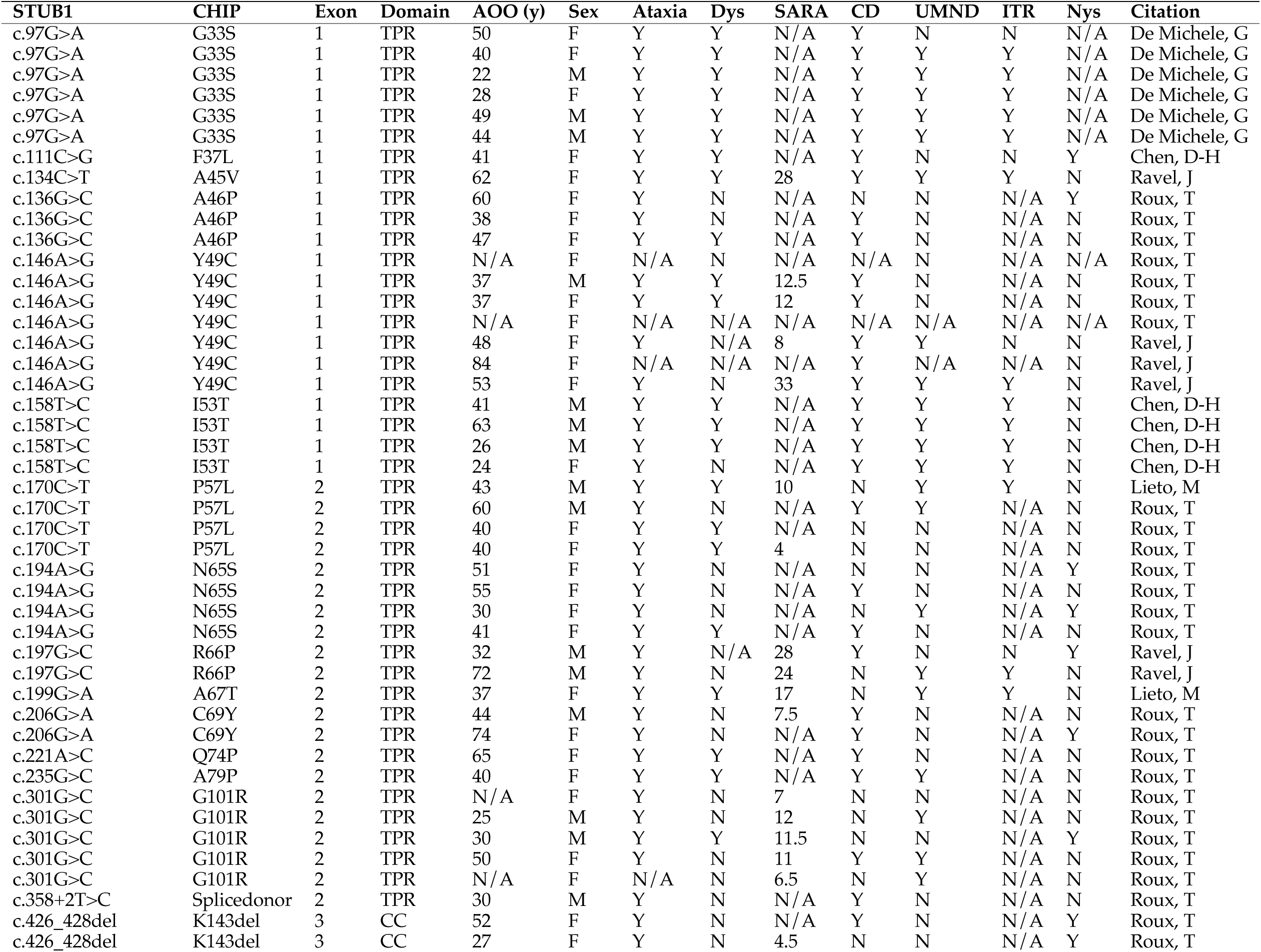
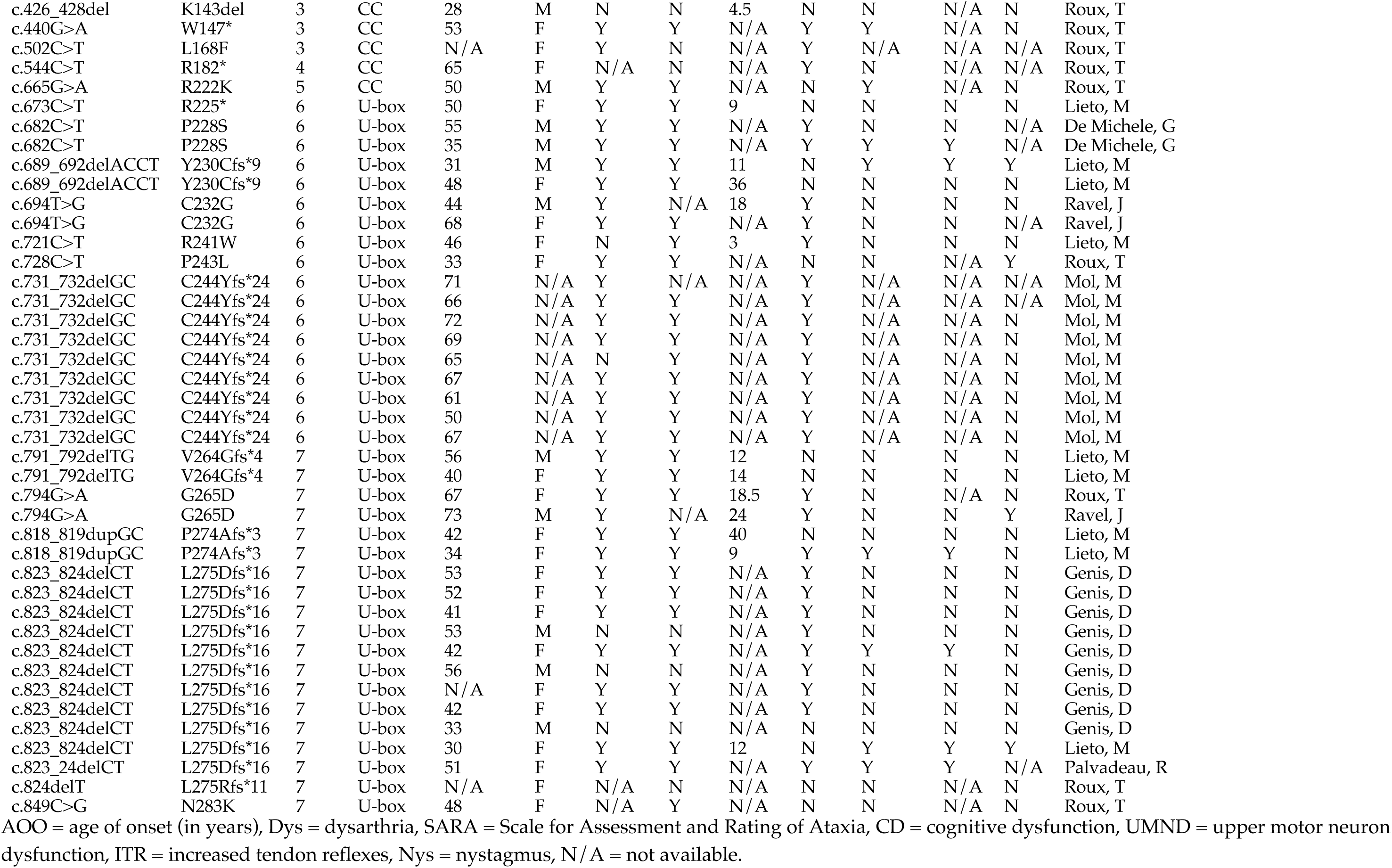
SCA48 Patient Genotype/Phenotype.

| STUB1 | CHIP | Exon | Domain | AOO (y) | Sex | Ataxia | Dys | SARA | CD | UMND | ITR | Nys | Citation |
| --- | --- | --- | --- | --- | --- | --- | --- | --- | --- | --- | --- | --- | --- |
| c.97G>A | G33S | 1 | TPR | 50 | F | Y | Y | N/A | Y | N | N | N/A | De Michele, G |
| c.97G>A | G33S | 1 | TPR | 40 | F | Y | Y | N/A | Y | Y | Y | N/A | De Michele, G |
| c.97G>A | G33S | 1 | TPR | 22 | M | Y | Y | N/A | Y | Y | Y | N/A | De Michele, G |
| c.97G>A | G33S | 1 | TPR | 28 | F | Y | Y | N/A | Y | Y | Y | N/A | De Michele, G |
| c.97G>A | G33S | 1 | TPR | 49 | M | Y | Y | N/A | Y | Y | Y | N/A | De Michele, G |
| c.97G>A | G33S | 1 | TPR | 44 | M | Y | Y | N/A | Y | Y | Y | N/A | De Michele, G |
| c.111C>G | F37L | 1 | TPR | 41 | F | Y | Y | N/A | Y | N | N | Y | Chen, D-H |
| c.134C>T | A45V | 1 | TPR | 62 | F | Y | Y | 28 | Y | Y | Y | N | Ravel, J |
| c.136G>C | A46P | 1 | TPR | 60 | F | Y | N | N/A | N | N | N/A | Y | Roux, T |
| c.136G>C | A46P | 1 | TPR | 38 | F | Y | N | N/A | Y | N | N/A | N | Roux, T |
| c.136G>C | A46P | 1 | TPR | 47 | F | Y | Y | N/A | Y | N | N/A | N | Roux, T |
| c.146A>G | Y49C | 1 | TPR | N/A | F | N/A | N | N/A | N/A | N | N/A | N/A | Roux, T |
| c.146A>G | Y49C | 1 | TPR | 37 | M | Y | Y | 12.5 | Y | N | N/A | N | Roux, T |
| c.146A>G | Y49C | 1 | TPR | 37 | F | Y | Y | 12 | Y | N | N/A | N | Roux, T |
| c.146A>G | Y49C | 1 | TPR | N/A | F | N/A | N/A | N/A | N/A | N/A | N/A | N/A | Roux, T |
| c.146A>G | Y49C | 1 | TPR | 48 | F | Y | N/A | 8 | Y | Y | N | N | Ravel, J |
| c.146A>G | Y49C | 1 | TPR | 84 | F | N/A | N/A | N/A | Y | N/A | N/A | N | Ravel, J |
| c.146A>G | Y49C | 1 | TPR | 53 | F | Y | N | 33 | Y | Y | Y | N | Ravel, J |
| c.158T>C | I53T | 1 | TPR | 41 | M | Y | Y | N/A | Y | Y | Y | N | Chen, D-H |
| c.158T>C | I53T | 1 | TPR | 63 | M | Y | Y | N/A | Y | Y | Y | N | Chen, D-H |
| c.158T>C | I53T | 1 | TPR | 26 | M | Y | Y | N/A | Y | Y | Y | N | Chen, D-H |
| c.158T>C | I53T | 1 | TPR | 24 | F | Y | N | N/A | Y | Y | Y | N | Chen, D-H |
| c.170C>T | P57L | 2 | TPR | 43 | M | Y | Y | 10 | N | Y | Y | N | Lieto, M |
| c.170C>T | P57L | 2 | TPR | 60 | M | Y | N | N/A | Y | Y | N/A | N | Roux, T |
| c.170C>T | P57L | 2 | TPR | 40 | F | Y | Y | N/A | N | N | N/A | N | Roux, T |
| c.170C>T | P57L | 2 | TPR | 40 | F | Y | Y | 4 | N | N | N/A | N | Roux, T |
| c.194A>G | N65S | 2 | TPR | 51 | F | Y | N | N/A | N | N | N/A | Y | Roux, T |
| c.194A>G | N65S | 2 | TPR | 55 | F | Y | N | N/A | Y | N | N/A | N | Roux, T |
| c.194A>G | N65S | 2 | TPR | 30 | F | Y | N | N/A | N | Y | N/A | Y | Roux, T |
| c.194A>G | N65S | 2 | TPR | 41 | F | Y | Y | N/A | Y | N | N/A | N | Roux, T |
| c.197G>C | R66P | 2 | TPR | 32 | M | Y | N/A | 28 | Y | N | N | Y | Ravel, J |
| c.197G>C | R66P | 2 | TPR | 72 | M | Y | N | 24 | N | Y | Y | N | Ravel, J |
| c.199G>A | A67T | 2 | TPR | 37 | F | Y | Y | 17 | N | Y | Y | N | Lieto, M |
| c.206G>A | C69Y | 2 | TPR | 44 | M | Y | N | 7.5 | Y | N | N/A | N | Roux, T |
| c.206G>A | C69Y | 2 | TPR | 74 | F | Y | N | N/A | Y | N | N/A | Y | Roux, T |
| c.221A>C | Q74P | 2 | TPR | 65 | F | Y | Y | N/A | Y | N | N/A | N | Roux, T |
| c.235G>C | A79P | 2 | TPR | 40 | F | Y | Y | N/A | Y | Y | N/A | N | Roux, T |
| c.301G>C | G101R | 2 | TPR | N/A | F | Y | N | 7 | N | N | N/A | N | Roux, T |
| c.301G>C | G101R | 2 | TPR | 25 | M | Y | N | 12 | N | Y | N/A | N | Roux, T |
| c.301G>C | G101R | 2 | TPR | 30 | M | Y | Y | 11.5 | N | N | N/A | Y | Roux, T |
| c.301G>C | G101R | 2 | TPR | 50 | F | Y | N | 11 | Y | Y | N/A | N | Roux, T |
| c.301G>C | G101R | 2 | TPR | N/A | F | N/A | N | 6.5 | N | Y | N/A | N | Roux, T |
| c.358+2T>C | Splicedonor | 2 | TPR | 30 | M | Y | N | N/A | Y | N | N/A | N | Roux, T |
| c.426_428del | K143del | 3 | CC | 52 | F | Y | N | N/A | Y | N | N/A | Y | Roux, T |
| c.426_428del | K143del | 3 | CC | 27 | F | Y | N | 4.5 | N | N | N/A | Y | Roux, T |
| c.426_428del | K143del | 3 | CC | 28 | M | N | N | 4.5 | N | N | N/A | N | Roux, T |
| c.440G>A | W147* | 3 | CC | 53 | F | Y | Y | N/A | Y | Y | N/A | N | Roux, T |
| c.502C>T | L168F | 3 | CC | N/A | F | Y | N | N/A | Y | N/A | N/A | N/A | Roux, T |
| c.544C>T | R182* | 4 | CC | 65 | F | N/A | N | N/A | Y | N | N/A | N/A | Roux, T |
| c.665G>A | R222K | 5 | CC | 50 | M | Y | Y | N/A | N | Y | N/A | N | Roux, T |
| c.673C>T | R225* | 6 | U-box | 50 | F | Y | Y | 9 | N | N | N | N | Lieto, M |
| c.682C>T | P228S | 6 | U-box | 55 | M | Y | Y | N/A | Y | N | N | N/A | De Michele, G |
| c.682C>T | P228S | 6 | U-box | 35 | M | Y | Y | N/A | Y | Y | Y | N/A | De Michele, G |
| c.689_692delACCT | Y230Cfs*9 | 6 | U-box | 31 | M | Y | Y | 11 | N | Y | Y | Y | Lieto, M |
| c.689_692delACCT | Y230Cfs*9 | 6 | U-box | 48 | F | Y | Y | 36 | N | N | N | N | Lieto, M |
| c.694T>G | C232G | 6 | U-box | 44 | M | Y | N/A | 18 | Y | N | N | N | Ravel, J |
| c.694T>G | C232G | 6 | U-box | 68 | F | Y | Y | N/A | Y | N | N | N/A | Ravel, J |
| c.721C>T | R241W | 6 | U-box | 46 | F | N | Y | 3 | Y | N | N | N | Lieto, M |
| c.728C>T | P243L | 6 | U-box | 33 | F | Y | Y | N/A | N | N | N/A | Y | Roux, T |
| c.731_732delGC | C244Yfs*24 | 6 | U-box | 71 | N/A | Y | N/A | N/A | Y | N/A | N/A | N/A | Mol, M |
| c.731_732delGC | C244Yfs*24 | 6 | U-box | 66 | N/A | Y | Y | N/A | Y | N/A | N/A | N/A | Mol, M |
| c.731_732delGC | C244Yfs*24 | 6 | U-box | 72 | N/A | Y | Y | N/A | Y | N/A | N/A | N | Mol, M |
| c.731_732delGC | C244Yfs*24 | 6 | U-box | 69 | N/A | Y | Y | N/A | Y | N/A | N/A | N | Mol, M |
| c.731_732delGC | C244Yfs*24 | 6 | U-box | 65 | N/A | N | Y | N/A | Y | N/A | N/A | N | Mol, M |
| c.731_732delGC | C244Yfs*24 | 6 | U-box | 67 | N/A | Y | Y | N/A | Y | N/A | N/A | N | Mol, M |
| c.731_732delGC | C244Yfs*24 | 6 | U-box | 61 | N/A | Y | Y | N/A | Y | N/A | N/A | N | Mol, M |
| c.731_732delGC | C244Yfs*24 | 6 | U-box | 50 | N/A | Y | Y | N/A | Y | N/A | N/A | N | Mol, M |
| c.731_732delGC | C244Yfs*24 | 6 | U-box | 67 | N/A | Y | Y | N/A | Y | N/A | N/A | N | Mol, M |
| c.791_792delTG | V264Gfs*4 | 7 | U-box | 56 | M | Y | Y | 12 | N | N | N | N | Lieto, M |
| c.791_792delTG | V264Gfs*4 | 7 | U-box | 40 | F | Y | Y | 14 | N | N | N | N | Lieto, M |
| c.794G>A | G265D | 7 | U-box | 67 | F | Y | Y | 18.5 | Y | N | N/A | N | Roux, T |
| c.794G>A | G265D | 7 | U-box | 73 | M | Y | N/A | 24 | Y | N | N | Y | Ravel, J |
| c.818_819dupGC | P274Afs*3 | 7 | U-box | 42 | F | Y | Y | 40 | N | N | N | N | Lieto, M |
| c.818_819dupGC | P274Afs*3 | 7 | U-box | 34 | F | Y | Y | 9 | Y | Y | Y | N | Lieto, M |
| c.823_824delCT | L275Dfs*16 | 7 | U-box | 53 | F | Y | Y | N/A | Y | N | N | N | Genis, D |
| c.823_824delCT | L275Dfs*16 | 7 | U-box | 52 | F | Y | Y | N/A | Y | N | N | N | Genis, D |
| c.823_824delCT | L275Dfs*16 | 7 | U-box | 41 | F | Y | Y | N/A | Y | N | N | N | Genis, D |
| c.823_824delCT | L275Dfs*16 | 7 | U-box | 53 | M | N | N | N/A | Y | N | N | N | Genis, D |
| c.823_824delCT | L275Dfs*16 | 7 | U-box | 42 | F | Y | Y | N/A | Y | Y | Y | N | Genis, D |
| c.823_824delCT | L275Dfs*16 | 7 | U-box | 56 | M | N | N | N/A | Y | N | N | N | Genis, D |
| c.823_824delCT | L275Dfs*16 | 7 | U-box | N/A | F | Y | Y | N/A | Y | N | N | N | Genis, D |
| c.823_824delCT | L275Dfs*16 | 7 | U-box | 42 | F | Y | Y | N/A | Y | N | N | N | Genis, D |
| c.823_824delCT | L275Dfs*16 | 7 | U-box | 33 | M | N | N | N/A | N | N | N | N | Genis, D |
| c.823_824delCT | L275Dfs*16 | 7 | U-box | 30 | F | Y | Y | 12 | N | Y | Y | Y | Lieto, M |
| c.823_24delCT | L275Dfs*16 | 7 | U-box | 51 | F | Y | Y | N/A | Y | Y | Y | N/A | Palvadeau, R |
| c.824delT | L275Rfs*11 | 7 | U-box | N/A | F | N/A | N | N/A | N | N | N/A | N | Roux, T |
| c.849C>G | N283K | 7 | U-box | 48 | F | N/A | Y | N/A | N | N | N/A | N | Roux, T |

AOO = age of onset (in years), Dys = dysarthria, SARA = Scale for Assessment and Rating of Ataxia, CD = cognitive dysfunction, UMND = upper motor neuron dysfunction, ITR = increased tendon reflexes, Nys = nystagmus, N/A = not available.

**Supplementary Table 3.**
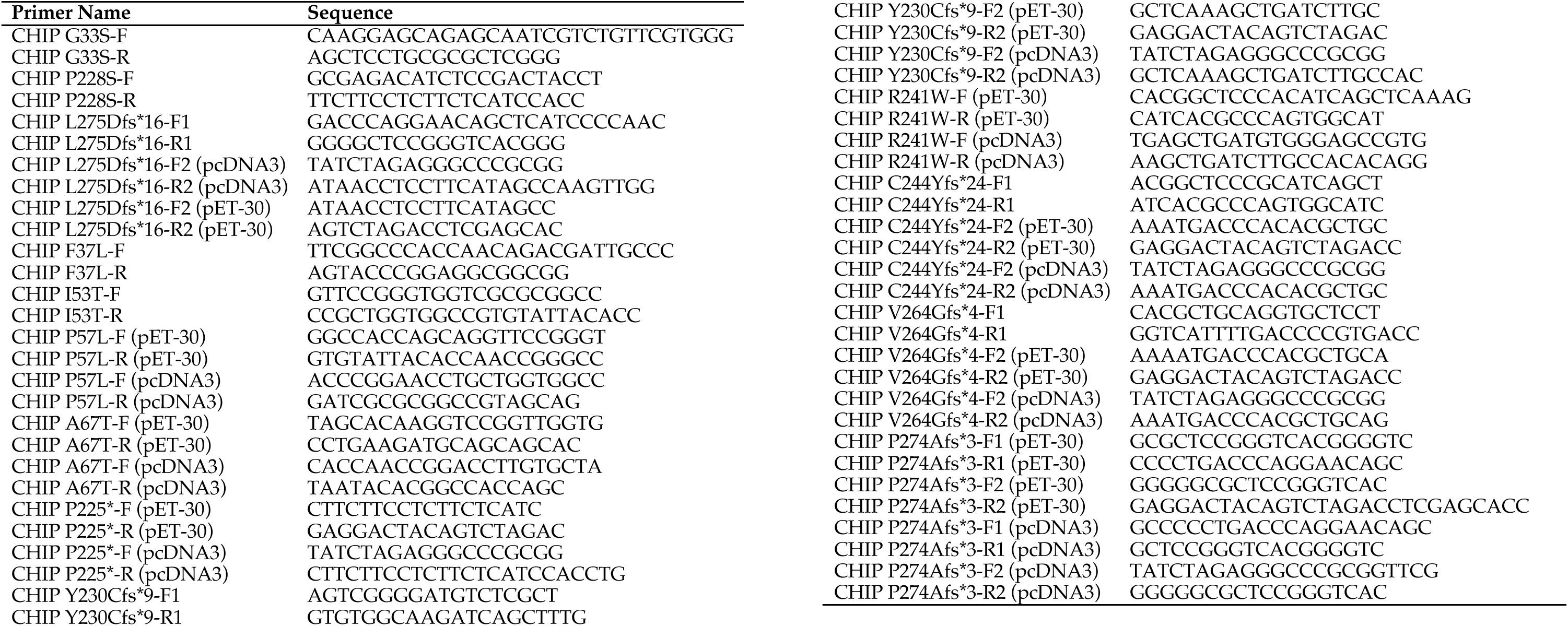
Primers used in CHIP Mutagenesis.

**Supplementary Table 4.** Primary antibodies used for immunoblotting and immunofluorescence.

| **Antibody** | **Source** | **Identifier** |
| --- | --- | --- |
| Myc-HRP | Cell Signaling Technologies | 2040 |
| Flag-HRP | Cell Signaling Technologies | 2044 |
| HA | Cell Signaling Technologies | 3724 |
| HSF1 | Enzo Life Sciences | ADI-SPA-901 |
| b-tubulin | Sigma | T3828 |
| Lamin B1-HRP | Cell Signaling Technologies | 15068 |
| Mouse Myc | Proteintech | 60003-2-Ig |
| PhosphoCHIP S20 | AbMart Inc.: p-CHIP Ser20* | 25011-1 |
| StarBright B700 | Bio-Rad | 12004161 |
| StarBright B520 | Bio-Rad | 12005866 |

\*(Custom antibody, Shanghai, China)

### Supplementary Methods

#### Nuclear and Cytoplasmic Fractionation

COS-7 cells were maintained in Dulbecco’s Modified Eagle’s Medium (DMEM; Corning, 10-017-CV) supplemented with 10% fetal bovine serum (FBS; Gibco, 26140-079) and 1% penicillin-streptomycin at 37 °C in a humidified 5% CO₂ incubator. For transient transfection, cells were seeded to reach ∼70–80% confluence and transfected with plasmid DNA using X-tremeGENE™ 9 DNA transfection reagent (Roche, 6365787001) at a 1:3 DNA-to-reagent ratio according to the manufacturer’s instructions. Approximately 24 hours after the trasnfection, nuclear and cytoplasmic extracts were prepared using the NE-PER Nuclear and Cytoplasmic Extraction Kit (Thermo Scientific, 78835) following the manufacturer’s protocol. Briefly, cells were harvested, lysed sequentially to separate cytoplasmic and nuclear components, and fractions were collected for downstream analyses.

### Supplementary Figure Legends

**Supplementary Figure 1.**
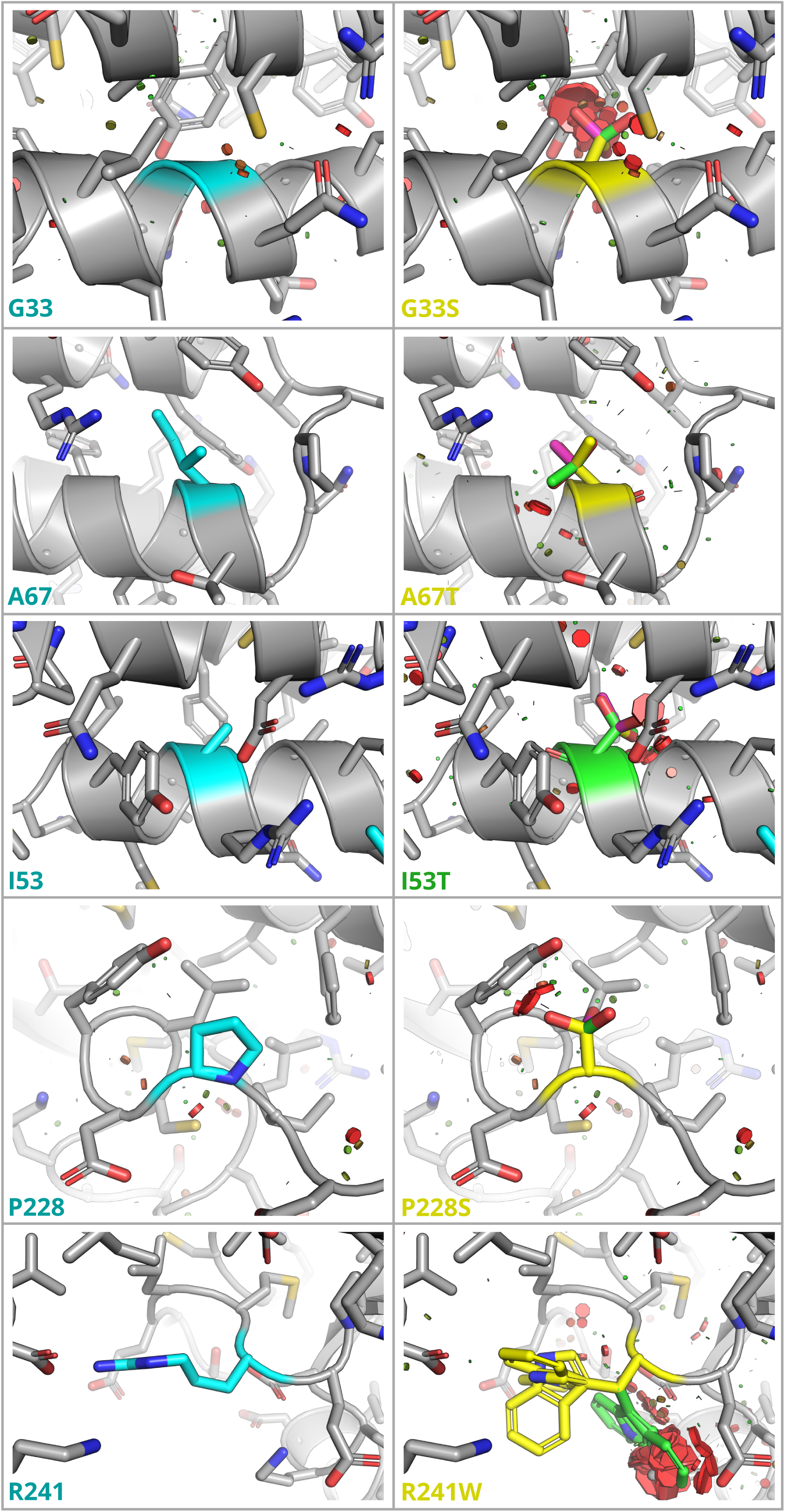
Structural modeling of CHIP variants reveals steric clashes at multiple mutation sites. The crystal structure of wild-type CHIP Wild-type (WT, grey) is shown as a cartoon representation with key side chains depicted as sticks. The native residues at positions 33 (glycine), 67 (alanine), 153 (isoleucine), 228 (proline), and 241 (arginine) are highlighted in cyan. Substitution of these residues with serine (G33S and P228S), threonine (A67T and I153T), or tryptophan (R241W) introduces new side-chain conformations (rotamers) shown as g⁺ (yellow), g⁻ (green), and t (magenta). Colored "bump" discs provide a qualitative indication of van der Waals interactions between mutant side chains and surrounding residues, binned as slight contact (green), slight overlap (orange), or significant overlap (red), with disc size proportional to the degree of steric interference.

**Supplementary Figure 2.**
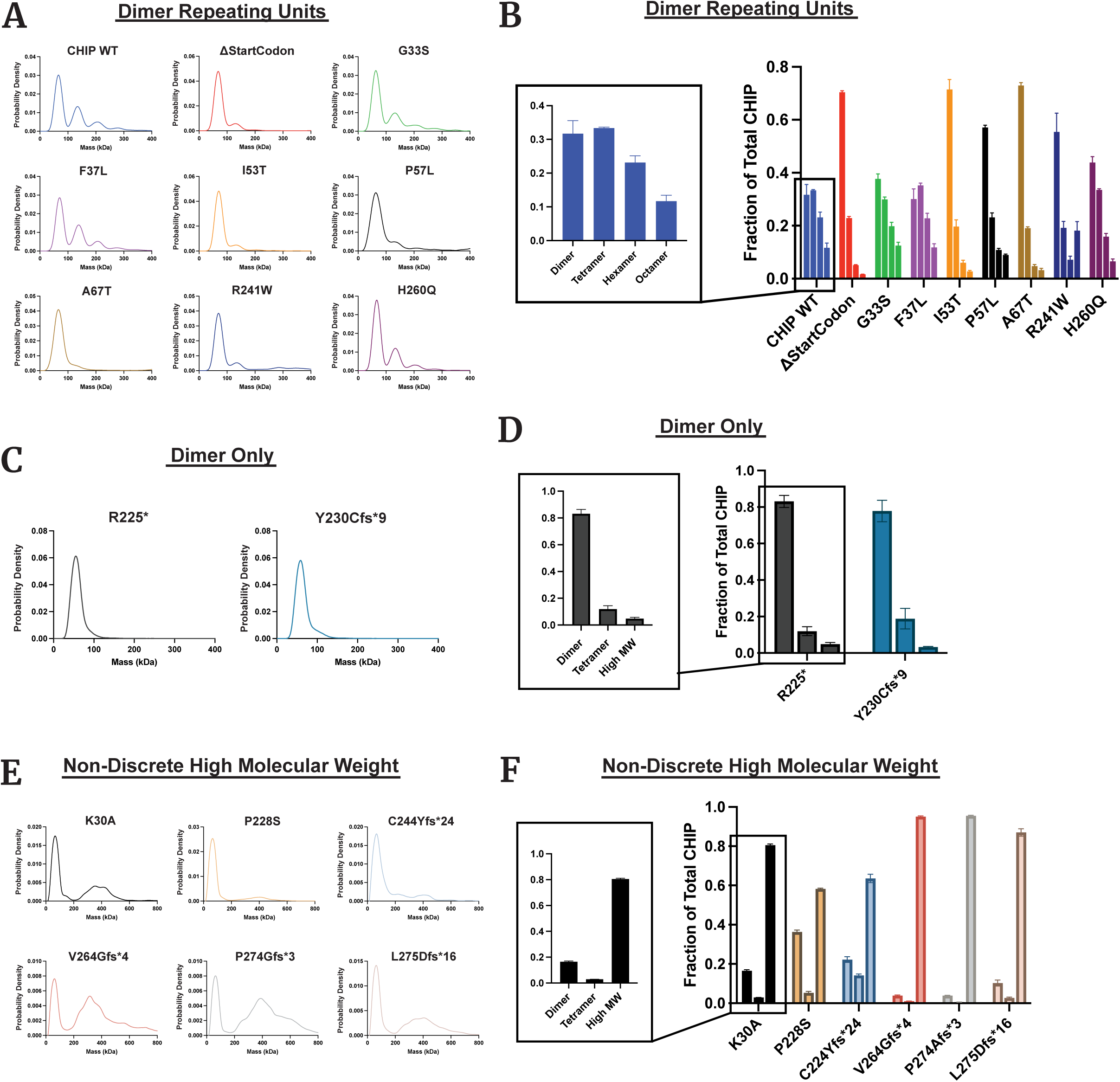
Mass photometry reveals distinct oligomerization profiles of CHIP wild type and SCA48-associated variants. We used mass photometry (MP) to monitor the oligomerization behavior of wild-type (WT) CHIP and SCA48-associated variants, based on the characteristic CHIP peaks corresponding to dimer, tetramer, hexamer, and octamer molecular weights. According to their MP profiles, CHIP mutants were grouped into three categories: *Dimer-repeating units:* (A) Representative MP profiles and (B) quantification of mutations ΔStartCodon, G33S, F37L, I53T, P57L, A67T, R241W, and H260Q, which retain distinct oligomeric states at the dimer, tetramer, hexamer, and octamer levels. *Dimer-only:* (C) MP profiles and (D) quantification for R225* and Y230Cfs*9* variants, which predominantly (∼80%) form dimers without detectable higher-order oligomers*. Non-discrete high molecular weight species: (E)* MP profiles and (F) quantification of synthetic mutations K30A, P228S, C244Yfs24, V264Gfs4, P274Gfs3, and L275Dfs*16, which exhibit very low levels of discrete oligomers and instead accumulate as non-discrete high-molecular-weight species.

**Supplementary Figure 3.**
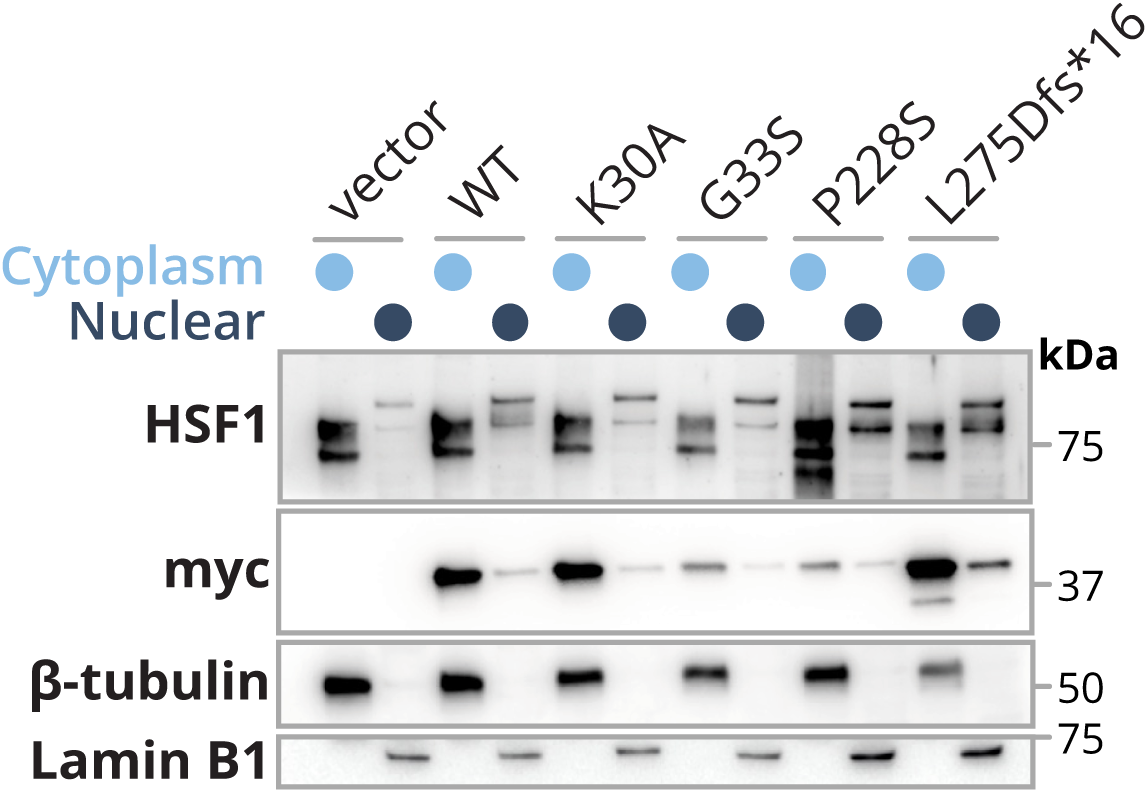
Subcellular localization of CHIP variants and HSF1 determined by fractionated immunoblotting and immunofluorescence. Fractionated immunoblotting of COS-7 cells expressing CHIP wild type (WT), G33S, P228S, L275Dfs16, and the synthetic mutant K30A revealed distinct nuclear and cytoplasmic distributions of CHIP and HSF1. CHIP WT was predominantly cytoplasmatic, but its overexpression enhanced nuclear localization of HSF1. The G33S, P228S, and K30A variants showed a mild reduction in nuclear CHIP compared to WT, whereas the L275Dfs16 mutant markedly increased nuclear accumulation of both CHIP and HSF1. CHIP WT overexpression increased HSF1 nuclear levels, which were generally retained—but not elevated—by most CHIP variants, with a remarkable increase observed for L275Dfs16.

**Supplementary Figure 4.**
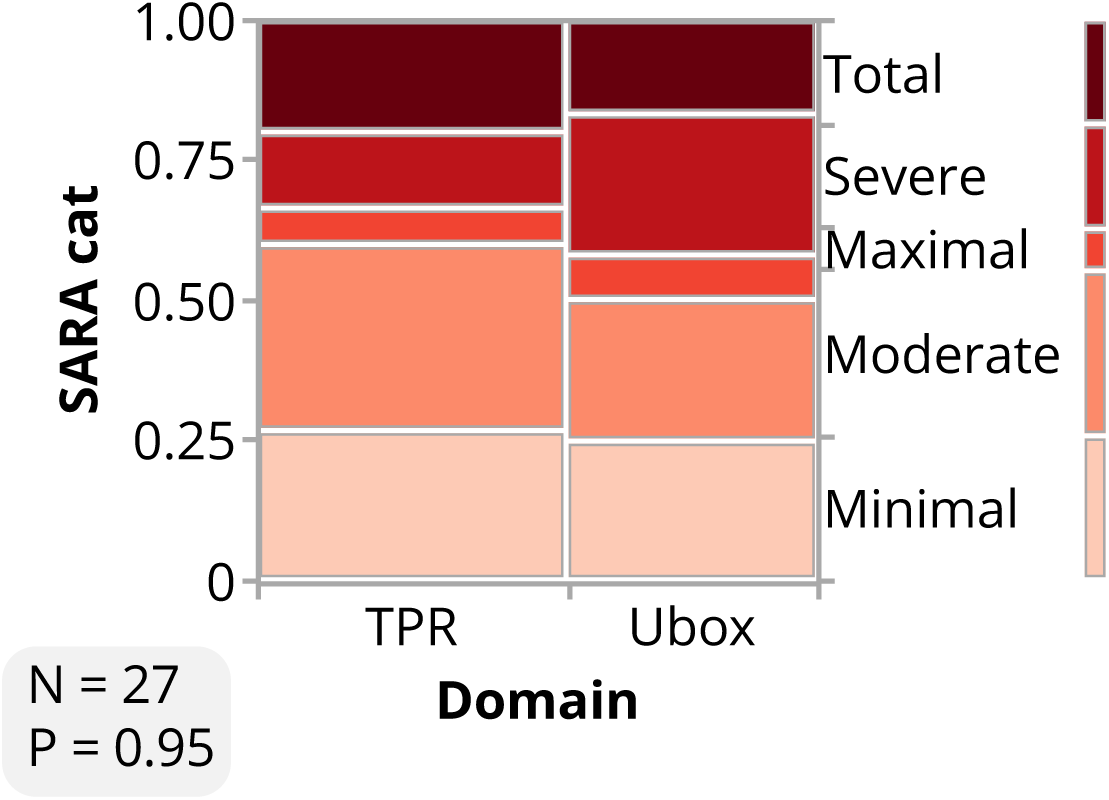
Relationship between CHIP mutation domain and clinical severity in SCA48 patients. Analysis of clinical data from 87 SCA48 patients with confirmed STUB1 mutations compiled from previously published case studies. The Scale for the Assessment and Rating of Ataxia (SARA) scores (when available, n=27) were plotted according to mutation location within the CHIP domains (TPR vs. U-box). Statistical comparison between the two groups was performed using an unpaired t-test. The number of data points and p-values are indicated in the bottom left corner of each plot. No significant difference in mean SARA scores was observed between patients carrying TPR or U-box domain mutations, suggesting that mutation domain alone does not account for disease severity variability.

